# High levels of cyclic diguanylate interfere with beneficial bacterial colonization

**DOI:** 10.1101/2022.04.11.487973

**Authors:** Ruth Y. Isenberg, David G. Christensen, Karen L. Visick, Mark J. Mandel

## Abstract

During colonization of the Hawaiian bobtail squid (*Euprymna scolopes*), *Vibrio fischeri* bacteria undergo a lifestyle transition from a planktonic motile state in the environment to a biofilm state in host mucus. Cyclic diguanylate (c-di-GMP) is a cytoplasmic signaling molecule that is important for regulating motility-biofilm transitions in many bacterial species. *V. fischeri* encodes 50 proteins predicted to synthesize and/or degrade c-di-GMP, but a role for c-di-GMP regulation during host colonization has not been investigated. We examined strains exhibiting either low or high levels of c-di-GMP during squid colonization and found that while a low c-di-GMP strain had no colonization defect, a high c-di-GMP strain was severely impaired. Expression of a heterologous c-di-GMP phosphodiesterase restored colonization, demonstrating that the effect is due to high c-di-GMP levels. In the constitutive high c-di-GMP state, colonizing *V. fischeri* exhibited reduced motility, altered biofilm aggregate morphology, and a regulatory interaction where transcription of one polysaccharide locus is inhibited by the presence of the other polysaccharide. Our results highlight the importance of proper c-di-GMP regulation during beneficial animal colonization, illustrate multiple pathways regulated by c-di-GMP in the host, and uncover an interplay of multiple exopolysaccharide systems in host-associated aggregates.

**IMPORTANCE:** There is substantial interest in studying cyclic diguanylate (c-di-GMP) in pathogenic and environmental bacteria, which has led to an accepted paradigm in which high c-di-GMP levels promote biofilm formation and reduce motility. However, considerably less focus has been placed on understanding how this compound contributes to beneficial colonization. Using the *Vibrio fischeri*-Hawaiian bobtail squid study system, we took advantage of recent genetic advances in the bacterium to modulate c-di-GMP levels and measure colonization and track c-di-GMP phenotypes in a symbiotic interaction. Studies in the animal host revealed a c-di-GMP-dependent genetic interaction between two distinct biofilm polysaccharides, Syp and cellulose, that was not evident in culture-based studies: elevated c-di-GMP altered the composition and abundance of the *in vivo* biofilm by decreasing *syp* transcription due to increased cellulose synthesis. This study reveals important parallels between pathogenic and beneficial colonization and additionally identifies c-di-GMP-dependent regulation that occurs specifically in the squid host.

## INTRODUCTION

In both pathogenic and beneficial associations, bacteria transition from a motile, planktonic state in the environment to a surface-associated biofilm state within the host. Cyclic diguanylate (c-di-GMP) is an intracellular signaling molecule that regulates this lifestyle transition for many bacterial species, including *Vibrio cholerae*, *Pseudomonas aeruginosa*, *Caulobacter crescentus*, and *Agrobacterium tumefaciens* (1, 2). Generally, higher intracellular c-di-GMP concentrations lead to increased biofilm formation and decreased motility, while lower intracellular c-di-GMP concentrations lead to the opposite phenotypes. C-di-GMP levels are regulated by diguanylate cyclase (DGC) proteins, which synthesize c-di-GMP, and phosphodiesterase (PDE) proteins, which degrade c-di-GMP. DGCs contain a GGDEF domain and PDEs contain either an EAL or HD-GYP domain. While the effects of c-di-GMP on motility, biofilm formation, and infectivity of pathogens such as *V. cholerae* have been extensively studied (3), the impact of c-di-GMP during the establishment of beneficial bacterial symbioses is not well understood. The focus of this study is to examine how c-di-GMP impacts beneficial colonization in a system where we can manipulate intracellular levels of the compound and dissect the colonization process in detail.

*Vibrio fischeri* is the exclusive, beneficial light organ symbiont of the Hawaiian bobtail squid (*Euprymna scolopes*). *V. fischeri* undergoes the motility-to-biofilm lifestyle transition during the initiation of host colonization (4–7). While *V. fischeri* is solitary and expresses polar flagella in seawater, upon encountering the juvenile bobtail squid light organ the bacteria require the symbiosis polysaccharide (Syp) to aggregate in the host mucus prior to commencing further colonization (5, 8–11). The bacterial aggregates form on the outer face of the light organ independent of flagellar motility and are influenced by physical currents generated by cilia on the light organ surface (4, 12). Host-produced nitric oxide stimulates dispersal from the biofilm aggregates (13), and the bacteria swim through the pore into the light organ ducts using flagellar motility and chemotaxis toward squid-derived chitin oligosaccharides (8, 14–17). The symbionts pass through an antechamber and bottleneck before reaching the crypts of the light organ, where they reach high density and produce light to camouflage the shadow of the squid host (18–20). The initiation of this process—including biofilm formation, chemotaxis, and motility—occurs only when the squid is newly hatched, as colonizing bacteria seed the host for its lifetime. Therefore, the mechanisms that regulate these processes are fundamental to selection for the correct symbionts and the establishment of the association.

As with the pathogens noted above, *V. fischeri* (strain ES114) encodes a large number of proteins predicted to regulate c-di-GMP levels: 50 total that encompass 28 DGCs, 14 PDEs, 5 DGC/PDEs, and 3 proteins that are predicted to be degenerate (21). Roles for DGCs MifA and MifB, and PDEs BinA and PdeV in *V. fischeri* motility and/or biofilm formation have been described (22–24). However, those studies did not examine individual mutants during squid colonization, and a large-scale transposon-insertion sequencing study did not identify individual mutations in genes related to c-di-GMP that were predicted to have significant phenotypes in the squid host (25). Given the predicted complexity of the c-di-GMP network in *V. fischeri*, we used new technology (26) to generate strains lacking multiple DGCs or PDEs to mimic either low or high c-di-GMP levels, and we applied these strains to determine the effects of c-di-GMP extremes on squid colonization. This approach allowed us to directly study the role of c-di-GMP *in vivo* during beneficial colonization, to determine bacterial behaviors that are influenced by the compound during animal colonization, and to reveal signaling interactions that are not evident during growth in culture.

## RESULTS

### Deletion of multiple DGCs or PDEs effectively manipulates c-di-GMP levels in the symbiont *V. fischeri*

To determine the role that c-di-GMP plays during the beneficial colonization of *E. scolopes* by *V. fischeri*, we constructed strains that were altered in their basal levels of c-di-GMP. While it is not yet feasible to delete all 50 c-di-GMP-related genes in a single strain, we applied recently developed technology that facilitates rapid construction of multiple gene deletions (26) to generate a strain that lacked seven DGCs (Δ7DGC; lacking *VF_1200*, *mifA*, *VF_1245*, *VF_A0342*-*A0343*, *VF_1639*, and *VF_A0216*) and another that lacked six PDEs (Δ6PDE; lacking *pdeV*, *VF_2480*, *binA*, *VF_0087*, *VF_1603*, and *VF_A0506*). These two strains were derived using a recently generated collection of 50 single mutants (27). The specific gene disruptions were chosen based both on technical reasons (*e.g.,* location of a gene of interest in the chromosome or the specific antibiotic resistance cassette present) and on preliminary phenotypic assessments. For the latter, we focused on mutations that, in single mutant experiments, impacted known c-di-GMP-dependent phenotypes, primarily the control over cellulose production (via Congo red binding) (24). Ultimately, our goal was to generate strains with decreased and increased levels of c-di-GMP. To determine if we achieved this goal, we examined the levels of c-di-GMP in these strains using the pFY4535 c-di-GMP reporter (28) and found that the Δ7DGC strain had c-di-GMP levels over 9-fold lower than the parent strain, while the Δ6PDE strain exhibited c-di-GMP levels about 1.5-fold higher (**Fig. 1A**). These differences were observed in rich medium as well as in filter-sterilized Instant Ocean (FSIO) under conditions that mimic squid inoculation (**Fig. 1A**), suggesting to us that these strains would be effective for studying the role of c-di-GMP during squid colonization. We examined phenotypes known to be regulated by c-di-GMP, including cellulose biofilm formation and flagellar motility. The Δ7DGC strain displayed elevated motility on TBS soft agar, reduced Congo red binding, and reduced activity of a *bcsQ’-gfp^+^* cellulose locus transcriptional reporter (**Fig. 1B-D**). The Δ6PDE strain exhibited the opposite phenotypes (**Fig. 1B-D**). These data supported our contention that we could use the multiple deletion approach to examine c-di-GMP regulation in the symbiont. Together, these data and those below support that the Δ7DGC strain represents cells in a low c-di-GMP state whereas the Δ6PDE cells are in a high c-di-GMP state, and for the remainder of the study we will refer to these strains as the “Low cdG” and “High cdG” strains, respectively.

**FIG 1.**
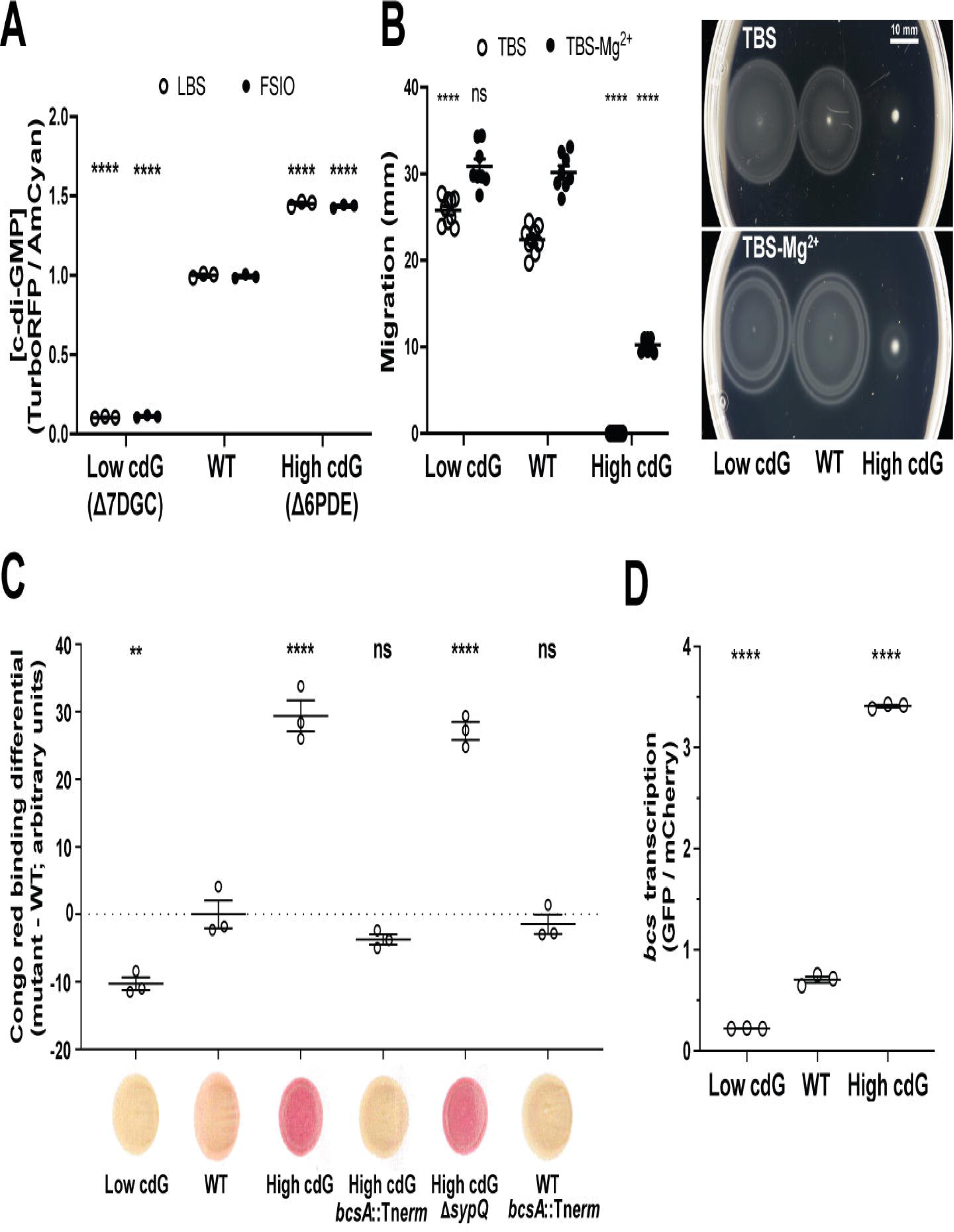
High c-di-GMP levels inhibit swimming motility, promote Congo red binding, and promote cellulose gene transcription. (A) Quantification of c-di-GMP concentration for indicated *V. fischeri* strains in LBS and filter-sterilized Instant Ocean (FSIO) using the pFY4535 c-di-GMP reporter plasmid. Constitutive AmCyan was used to normalize GFP to cell density. For each strain, *n* = 3 biological and *n* = 3 technical replicates per biological replicate, each point represents the mean of technical replicates, average bars represent the mean of biological replicates, and error bars represent standard error of the mean. (B) Quantification of migration through soft (0.3%) agar for indicated *V. fischeri* strains. For each strain, *n* = 8-9 biological replicates, average bars represent mean, and error bars represent standard error of the mean. Motility plate images are representative. (C) Quantification of Congo red binding for indicated *V. fischeri* strains. For each strain, *n* = 3 technical replicates, average bars represent mean, and error bars represent standard error of the mean. Congo red spot images are representative. (D) Quantification of *in vitro bcs* transcription by indicated *V. fischeri* strains using the pRYI063 *bcsQ*’-*gfp*^+^ transcriptional reporter plasmid. For each strain, n = 3 biological replicates. Points represent each biological replicate, average bars represent the mean, and error bars represent standard error of the mean. Constitutive mCherry was used to normalize GFP to cell density. For panels A-D, one-way ANOVA was used for statistical analysis and asterisks represent significance relative to the WT strain; ns = not significant, *p < 0.05, **p < 0.006, ****p < 0.0001.

### Elevated c-di-GMP levels interfere with productive colonization initiation

To determine whether the lower and/or higher c-di-GMP levels impacted symbiotic colonization, we introduced the Low cdG and High cdG strains to newly hatched *E. scolopes* squid. Each strain was individually inoculated into a bowl with hatchling squid and allowed to initiate colonization for 3 h. The squid were transferred to new water, and the bacterial load in each host was measured at 18 and 48 hours-post-inoculation (hpi) (29). For WT *V. fischeri* ES114, this process leads to approximately 10^5^ CFU per squid at 18 and 48 hpi, and a similar yield was observed for the Low cdG strain (**Fig. 2**). This was surprising to us, given that there are 50 genes that encode c-di-GMP synthesis/degradation factors, yet downregulation of c-di-GMP levels by over 9-fold did not impact the ability to colonize the squid host (**Fig. 2**). To ask whether the low cdG strain exhibited a more subtle colonization defect that we were not detecting in this single strain colonization assay, we performed a competitive colonization assay in which the Low cdG and WT strain were co-inoculated. We reasoned that if the Low cdG strain had a slight defect in the host, then the competitive assay could reveal this effect. However, in the competitive assay, the Low cdG strain exhibited similar fitness as the WT strain (**Fig. S1**). In contrast, in the single-strain colonization assay, the High cdG strain had a substantial deficit in bacterial yield in the host as early as 18 hpi and also at 48 hpi, with a median CFU below 10^3^ per light organ (**Fig. 2**). In the competitive colonization, the High cdG strain displayed a significant deficit, as WT outcompeted the High cdG strain by over 3000-fold (**Fig. S1**). Growth of the High cdG strain in culture was similar to WT (**Fig. S2**), arguing that the colonization defect is due to a developmental deficit in the host and not an inherent growth property of the strain.

**FIG 2.**
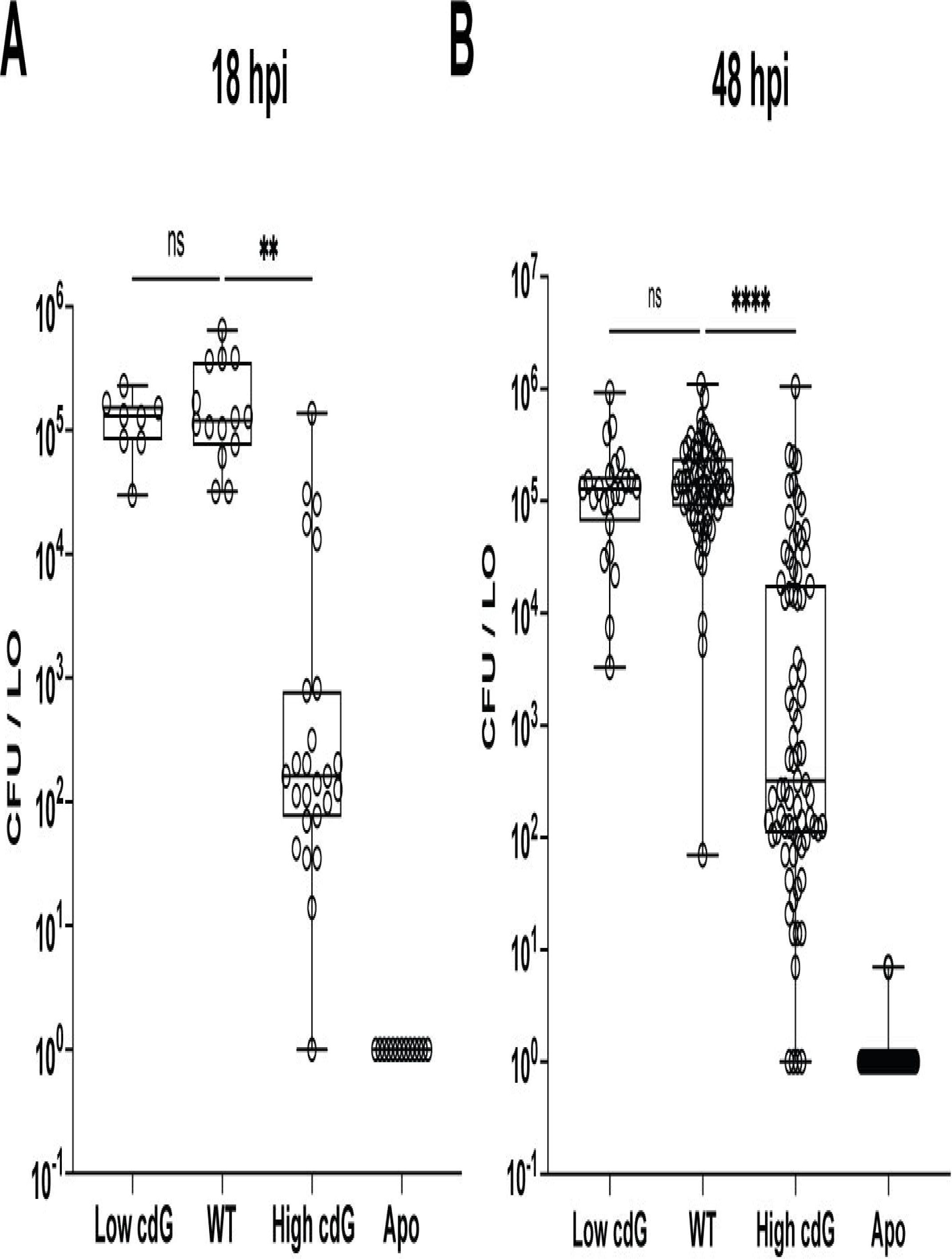
Elevated c-di-GMP levels inhibit host colonization. (A) Quantification of squid colonization levels at 18 hpi by indicated *V. fischeri* strains and aposymbiotic (Apo) control. Sample sizes from left to right are 8, 14, 25, and 13 squid. (B) Quantification of squid colonization levels at 48 hpi by indicated *V. fischeri* strains and aposymbiotic (Apo) control. Sample sizes from left to right are 23, 68, 75, and 65 squid. For panels A and B, box-and-whisker plots represent minimum, 25th percentile, median, 75th percentile, and maximum. Kruskal-Wallis test was performed for statistical analysis for squid that were introduced to bacteria; ns = not significant,**p < 0.002, ****p < 0.0001.

From these data, we conclude that high levels of c-di-GMP are detrimental for the initiation of a beneficial colonization in the squid light organ, whereas low levels of c-di-GMP do not impact the colonization behavior during the first 48 h of symbiosis.

To ask whether the phenotypes we observed in the High cdG strain were due to regulation of c-di-GMP levels or whether they were due to a specific gene product that is absent (i.e., signaling specificity), we introduced a PDE from *V. cholerae*, VC1086, that is known to be active in heterologous bacteria (30, 31), and that we predicted would lower c-di-GMP levels in *V. fischeri*. Across the phenotypes tested, the plasmid-expressed VC1086 had only a slight effect in a WT background (**Fig. 3**). However, in the High cdG background, expression of VC1086 reduced cdG levels by 1.6-fold, restored swimming motility on TBS agar, and reduced Congo red binding 1.7-fold in the High cdG background (**Fig. 3A-C**). In a squid colonization assay, expression of the *V. cholerae* PDE rescued colonization of the High cdG strain (**Fig. 3D**). This set of experiments confirmed that our results were due to altering levels of c-di-GMP rather than the absence of a single gene product. We conclude that lower levels of c-di-GMP are not harmful to establishing a symbiotic association, while higher levels are detrimental to symbiotic initiation.

**FIG 3.**
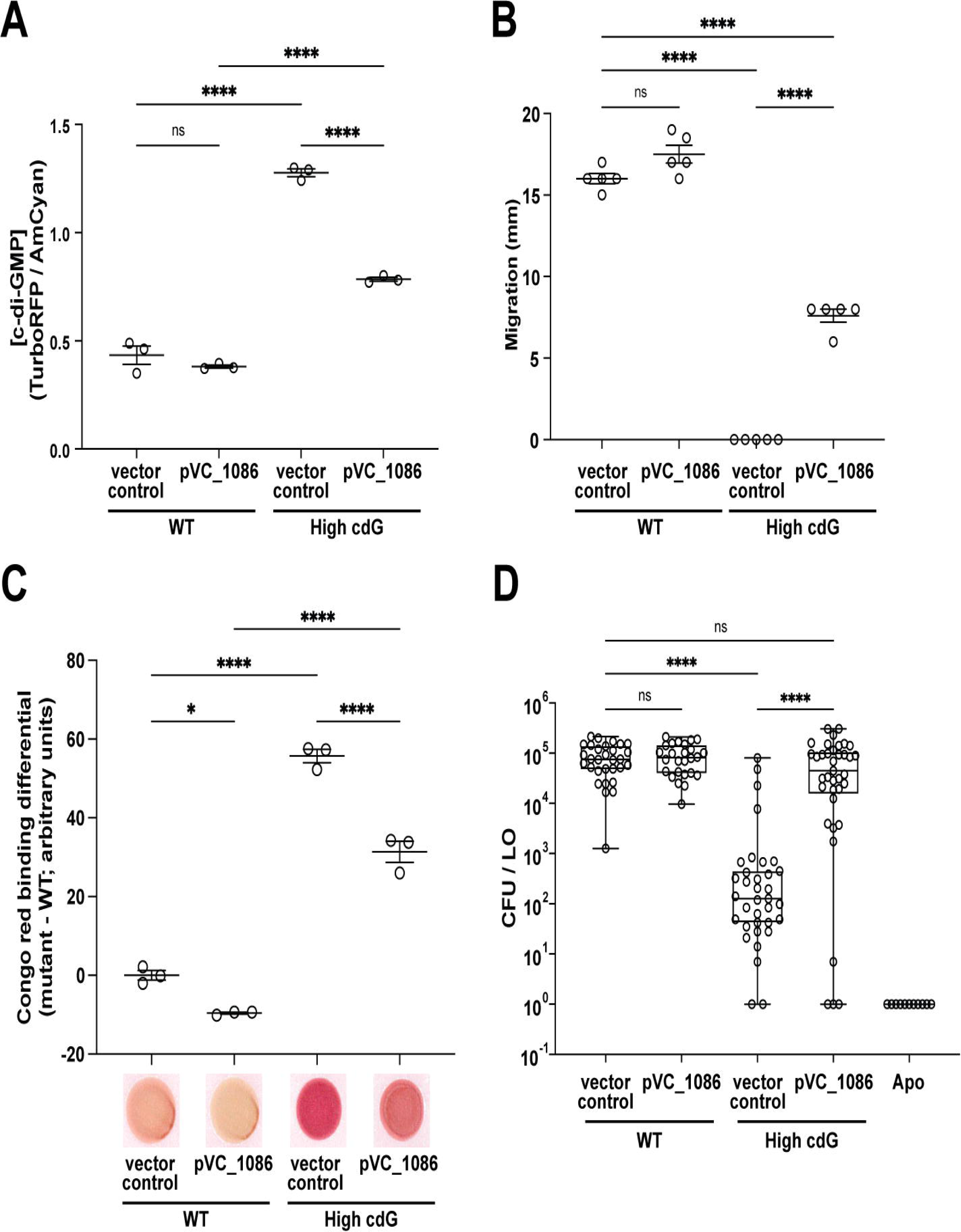
Reduced swimming motility, increased Congo red binding, and diminished squid colonization of the High cdG strain are dependent on high c-di-GMP levels. (A) Quantification of c-di-GMP concentration for indicated *V. fischeri* strains using the pFY4535 c-di-GMP reporter plasmid. For each strain, *n* = 3 biological and *n* = 3 technical replicates, each dot represents the mean of technical replicates, average bars represent the mean of biological replicates, and error bars represent standard error of the mean. (B) Quantification of migration through TBS soft (0.3%) agar for indicated *V. fischeri* strains. For each strain, *n* = 5 biological replicates, average bars represent mean, and error bars represent standard error of the mean. (C) Quantification of Congo red binding for indicated *V. fischeri* strains. For each strain, *n* = 3 technical replicates, average bars represent the mean, and error bars represent standard error of the mean. Congo red spot images are representative. For panels A-C, one-way ANOVA was used for statistical analysis; ns = not significant, *p = 0.017, ****p < 0.0001. (D) Quantification of squid colonization levels at 48 hpi by indicated *V. fischeri* strains and aposymbiotic (Apo) control. Box-and-whisker plots represent minimum, 25th percentile, median, 75th percentile, and maximum. Sample sizes from left to right are 30, 26, 34, 37, and 11 squid. Kruskal-Wallis test was performed for statistical analysis for squid that were introduced to bacteria; ns = not significant, ****p < 0.0001.

### The High c-di-GMP strain exhibits flagellar motility *in vivo*

In culture, the High cdG strain had a substantial swimming motility defect (**Fig. 1B**). We therefore considered the hypothesis that a key basis for the poor colonization by the High cdG strain was due to the altered flagellar motility, which is required for host colonization (8, 15, 17). In culture, we did observe a migration ring in the presence of magnesium sulfate, which promotes *V. fischeri* motility (32), demonstrating that the strain is not amotile (**Fig. 1B**). Cells from the outer ring of motility from the High cdG strain (**Fig. 1B**) exhibited the same phenotype as the parent High cdG strain upon reinoculation into a new plate (i.e., low motility) (**Fig. S3**), arguing that the motility observed was not due to spontaneous suppressor mutants. An open question, then, was whether this strain exhibited flagellar motility under the conditions experienced in the light organ mucus. We therefore performed a competitive colonization assay between the High cdG strain and the amotile High cdG Δ*flrA* isogenic derivative. The High cdG strain outcompeted the amotile Δ*flrA* derivative, with a median advantage of approximately 10-fold (**Fig. 4**), supporting that the High cdG strain is indeed motile in the host.

**FIG 4.**
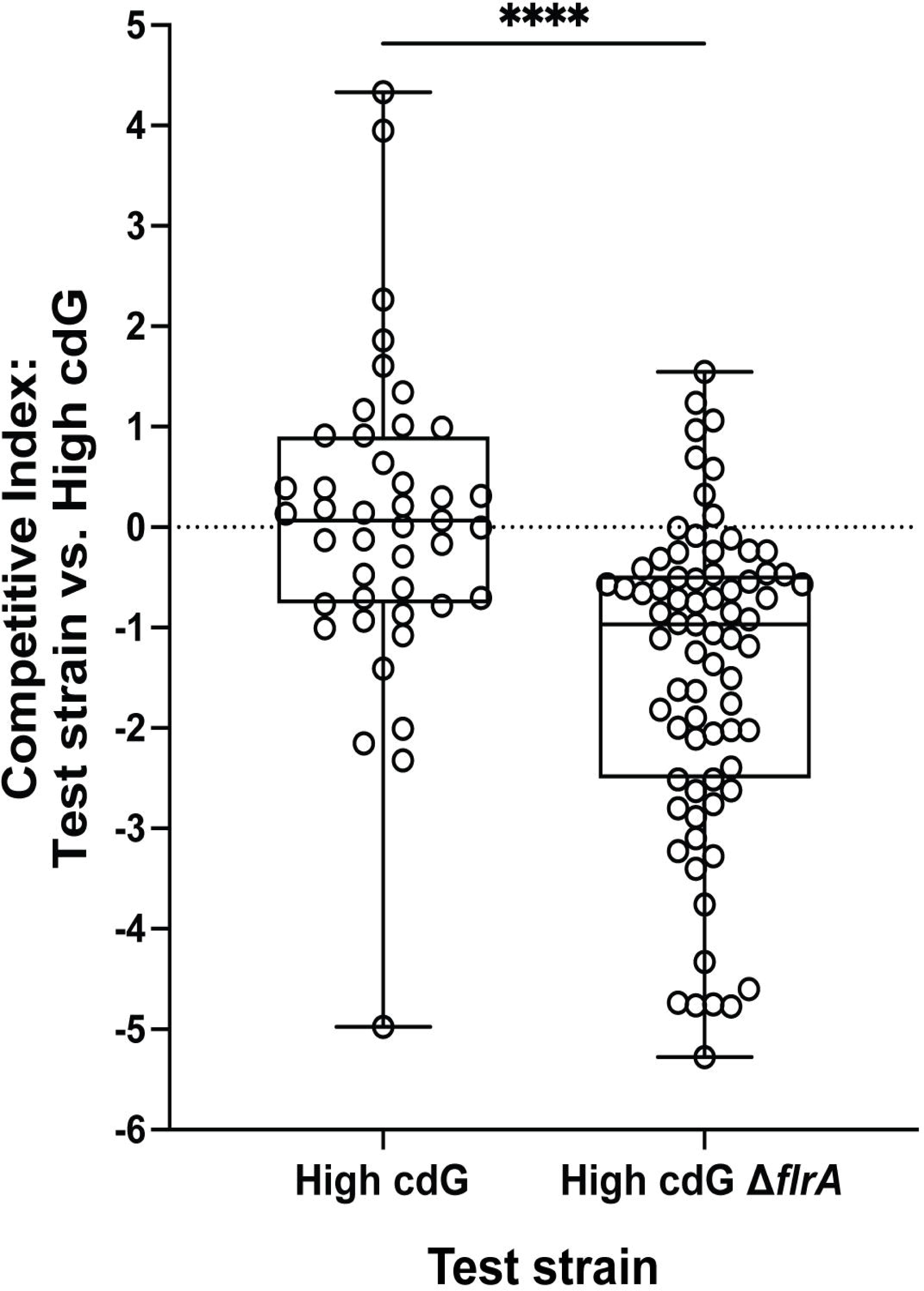
High c-di-GMP levels do not eliminate motility during host colonization. Quantification of squid competitive colonization index at 48 hpi by indicated *V. fischeri* strains. Competitive index represents log_10_((test strain/High cdG)_output_ / (test strain/High cdG)_input_). Box-and-whisker plots represent minimum, 25th percentile, median, 75th percentile, and maximum. Sample sizes from left to right are 43 and 75 squid. Kruskal-Wallis test was performed for statistical analysis; ****p < 0.0001.

### Elevated c-di-GMP levels lead to altered aggregate morphology *in vivo*

The *V. fischeri*-squid system provides an opportunity to combine genetic and imaging approaches to study the planktonic-to-biofilm transition during colonization of a symbiotic host. Symbiosis polysaccharide (Syp) is required for host colonization (5), while the distinct cellulose polysaccharide is not known to impact host phenotypes, so we questioned whether c-di-GMP would impact Syp-dependent aggregation *in vivo*. At 3 hpi, we examined *in vivo* biofilm aggregates using *V. fischeri* that constitutively expressed a fluorescent protein (GFP or mCherry). Compared to the Low cdG strain, the High cdG strain produced larger-sized aggregates, while the WT aggregates were an intermediate size (**Fig. 5A,B**). In over 10% of the light organs examined, the High cdG strain produced four or more aggregates, whereas squid colonized by the WT strain had three aggregates at most (**Fig. 5C**).

**FIG 5.**
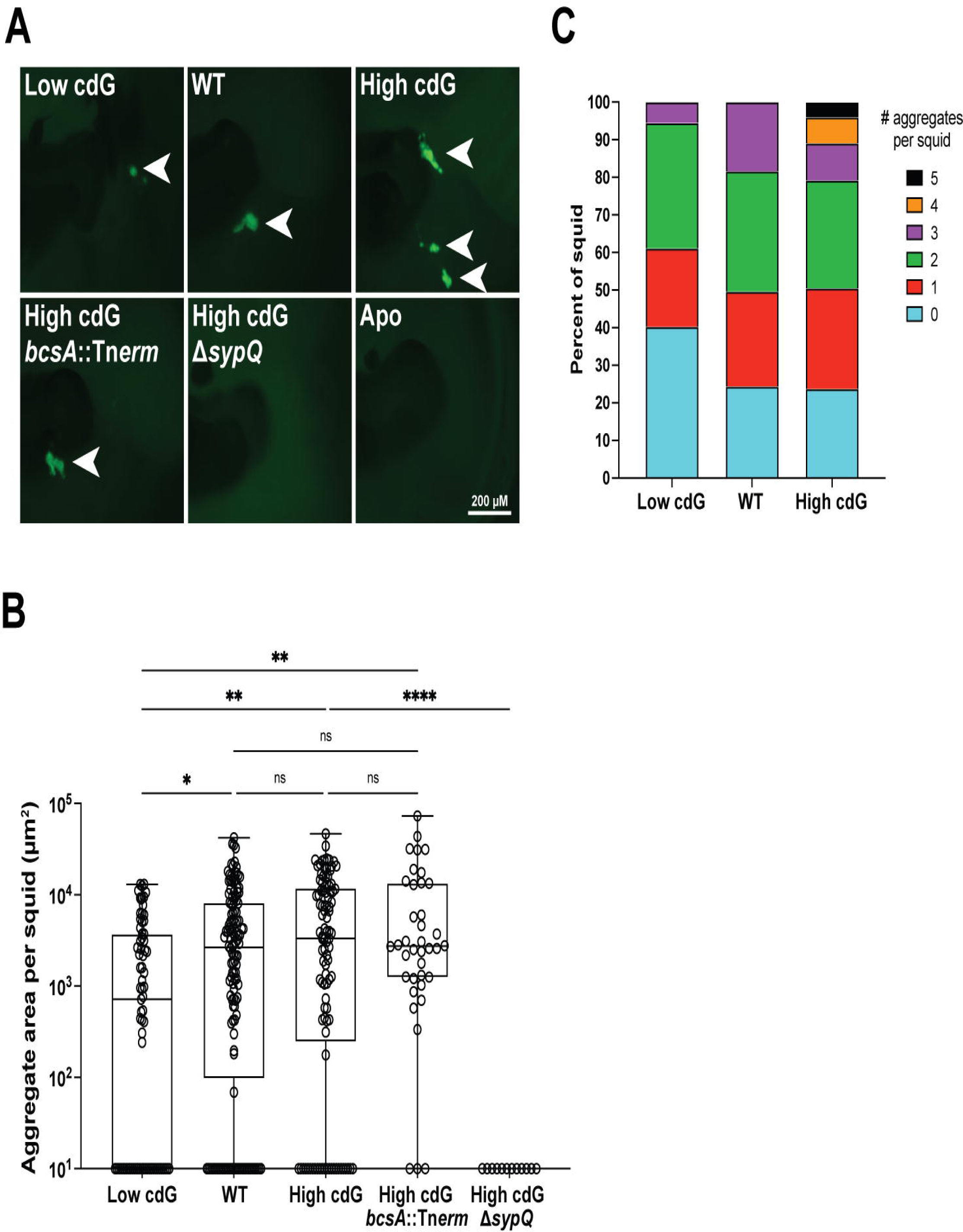
Elevated c-di-GMP levels lead to a greater number of *V. fischeri* aggregates in the squid host. (A) Representative fluorescence microscopy images of aggregates made by indicated *V. fischeri* strains carrying a constitutive GFP on the pVSV102 plasmid within the host mucus. Arrows indicate location of aggregates. (B) Quantification of aggregate area per squid for indicated *V. fischeri* strains carrying a constitutive GFP on the pVSV102 plasmid. Zero values are displayed at the bottom of the plot. Box-and-whisker plots represent minimum, 25th percentile, median, 75th percentile, and maximum. Sample sizes from left to right are 73, 131, 101, 38, and 12 squid. Kruskal-Wallis test was used for statistical analysis; ns = not significant, *p < 0.04, **p < 0.02, ****p < 0.0001. (C) Quantification of number of aggregates formed per squid by indicated *V. fischeri* strains. Sample sizes from left to right are 73, 131, and 101 squid.

Although the pFY4535 reporter plasmid was not designed for *in vivo* squid experiments, we attempted to use this tool to measure c-di-GMP levels in the host for the Low cdG, WT, or High cdG strain (28). AmCyan is produced constitutively from the plasmid, enabling us to locate the bacterial cells in the host. Turbo RFP as a reporter for c-di-GMP levels was substantially reduced in the Low cdG strain compared to WT, consistent with what we observed *in vitro* (**Fig. S4, Fig. 1**). Examination of the High cdG strain, however, revealed aggregate morphology that was not consistent with what we observed above with a constitutive GFP marking the cells in aggregates (**Fig. 5**). Specifically, we saw diffuse small groups of cells in the host in the High cdG/pFY4535 strain, rather than the aggregates that are produced by the High cdG/pVSV102 (GFP) strain. The High cdG/pFY4535 phenotype is the outlier, as the gene reporter assays described in the next paragraph use a constitutive mCherry to mark the cells, and their aggregate behavior matches that of the constitutive GFP in **Fig. 5**. Put together, our interpretation of these data is that the high level of reporter expression in the High cdG strain *in vivo* alters the aggregative behavior of the cells. While this effect prohibits accurate measurements of c-di-GMP levels in the host, it is consistent with a model in which high c-di-GMP levels in the strain are retained *in vivo*.

To test if the Syp polysaccharide was required for the pattern of multiple bacterial aggregates we observed in the High cdG strain, we deleted the gene encoding the structural Syp protein SypQ in the High cdG background. This strain failed to colonize the host robustly at 48 hpi (**Fig. S5B**) and did not form detectable aggregates (**Fig. 5A,B**), demonstrating that Syp-dependent biofilm is required for *in vivo* aggregation in the High cdG background. We asked whether transcription of the *syp* locus was altered in the High cdG strain. Activity of a *sypA’-gfp^+^*reporter was examined during the aggregation phase, where we found that in the host, *sypA’-gfp^+^* reporter activity was substantially reduced in the High cdG strain compared to the WT parent (**Fig. 6A, Fig. S6A**). High levels of c-di-GMP therefore negatively impact the expression of the symbiosis polysaccharide locus *in vivo*. Notably, when we asked whether the High cdG strain had altered *sypA’-gfp^+^* reporter activity *in vitro*, we found no significant effects in LBS medium or in TBS-Ca^2+^ medium that leads to a higher basal *syp* level (**Fig. 6C**) (33). Therefore, c-di-GMP exerts a host-specific effect on symbiotic biofilm expression.

**FIG 6.**
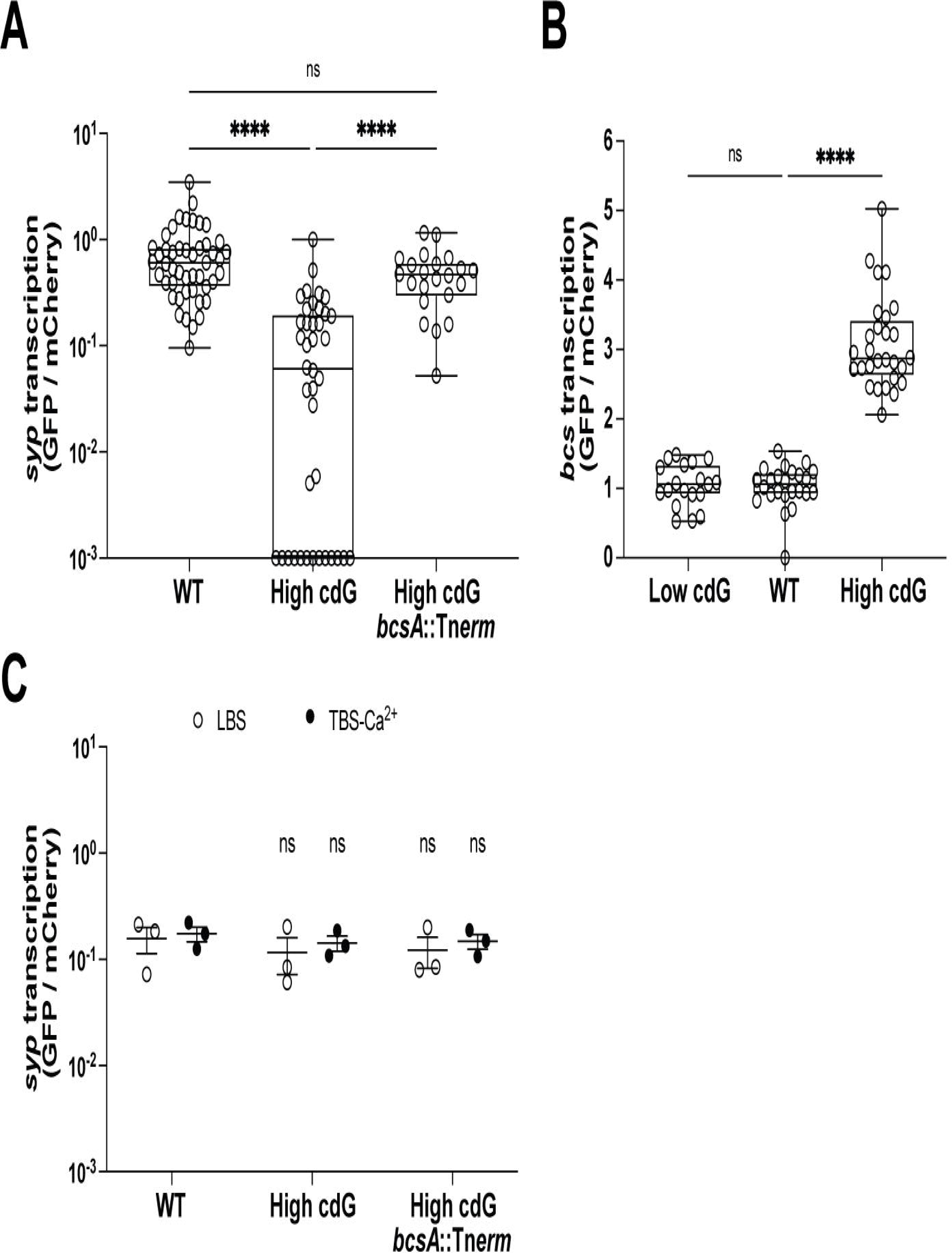
c-di-GMP downregulates *syp* via BcsA in host mucus but not in culture. (A) Quantification of *syp* transcription for indicated *V. fischeri* strains using the pM1422 *sypA*’-*gfp*^+^ transcriptional reporter plasmid in aggregates within the host mucus. Sample sizes from left to right are 47, 40, and 22 aggregates. (B) Quantification of *bcs* transcription for indicated *V. fischeri* strains using the pRYI063 *bcsQ*’-*gfp*^+^ transcriptional reporter plasmid in aggregates within the host mucus. Sample sizes from left to right are 19, 24 and 28 aggregates. For panels A and B, Box-and-whisker plots represent minimum, 25th percentile, median, 75th percentile, and maximum. Kruskal-Wallis test was used for statistical analysis; ns = not significant, ****p < 0.0001. (C) Quantification of *in vitro syp* transcription in LBS and TBS-Ca^2+^ for indicated *V. fischeri* strains carrying the pM1422 *sypA*’*-gfp*^+^ transcriptional reporter plasmid. For each strain, *n* = 3 biological replicates. Points represent each replicate, average bars represent the mean of replicates, and error bars represent the standard error of the mean. One-way ANOVA was used for statistical analysis; ns = not significant relative to the WT strain in the same media. For panels A-C, constitutive mCherry was used to normalize GFP to cell density.

### High levels of c-di-GMP repress *syp* polysaccharide transcription via the cellulose polysaccharide and in a host-dependent manner

In previous studies in the squid host, larger bacterial aggregates have correlated with an upregulation of *syp* gene transcription and a higher propensity for colonization initiation and competitive fitness (5, 34, 35). In this case, however, the larger aggregates of the High cdG strain have reduced *sypA’-gfp^+^* reporter activity and exhibit poor colonization (**Fig. 6A, Fig. S6A, Fig. 2**). We therefore asked if there was another component of the aggregate that may contribute to the morphology observed but that may not enhance colonization ability in the same way that additional Syp polysaccharide promotes colonization. A candidate for this component is cellulose, as we demonstrated above that cellulose is upregulated in the High cdG strain (**Fig. 1C,D**). Therefore, we asked whether there may be an interaction between the cellulose and Syp polysaccharide biosynthetic pathways. We measured *bcsQ* transcription *in vivo* using a fluorescent transcriptional reporter (*bcsQ’-gfp^+^*), and we determined that *bcsQ’-gfp^+^* reporter activity was elevated in bacterial aggregates in the host mucus in the High cdG strain compared to WT (**Fig. 6B, Fig. S6B**). Using a *bcsA*::Tn*erm* transposon insertion mutant that interrupts cellulose synthase, we examined colonization in the context of the High cdG background.

Interruption of *bcsA* did not improve colonization at 18 or 48 hpi compared to the High cdG strain (**Fig. S5**). However, compared to the High cdG strain, a *bcsA* derivative exhibited higher *sypA’-gfp^+^*activity *in vivo* (**Fig. 6B, Fig. S6B**), yet no difference in *sypA’-gfp*^+^ activity was observed *in vitro* with the same strains, even with *syp* induced by added calcium chloride (**Fig. 6C**) (33). These data present a novel connection between transcriptional control of polysaccharide systems that occurs specifically in the host.

## DISCUSSION

### Cyclic di-GMP impacts multiple pathways during beneficial colonization

Our study was motivated by the question of how c-di-GMP impacts a beneficial microbe-host association. To ask this question, we focused on the *V. fischeri*-squid mutualism. This well-studied binary association allows us to examine individual stages of colonization using strains deleted for 6-7 genes and that consequently have altered levels of c-di-GMP. We found that a strain with elevated levels of c-di-GMP exhibited striking defects during host colonization. The High cdG strain formed aggregates that trended larger (**Fig. 4**), had altered host-specific polysaccharide gene expression in the aggregates (**Fig. 6A,B, Fig. S6**), exhibited reduced flagellar motility (**Fig. 1B**), and displayed a significant colonization defect as enumerated by bacterial counts in the host as early as 18 hpi (**Fig. 2**). Bacterial counts did not recover by 48 hpi, suggesting that high levels of c-di-GMP confer both an initiation defect and an accommodation defect on the colonizing strains (36), as discussed further below. Colonization of the High cdG strain was rescued by overexpression of a heterologous PDE that does not have significant homology to proteins in *V. fischeri* (**Fig. 3D**), strongly supporting that the observed effects are due to the c-di-GMP levels and not to individual functions of the gene products affected in the multiple deletion strains.

Our results parallel work in several pathogenic bacteria, where elevated c-di-GMP levels are correlated with a decrease in virulence (3, 37–40). In *Yersinia pestis*, the small number of relevant enzymes made it possible to remove all of the DGCs or PDEs. Deletion of both genes for the functional DGCs effectively eliminated c-di-GMP in the cell (measured by LC-MS), yet virulence in mouse infection models was not affected (40). However, deletion of the single functional PDE gene, *hmsP*, led to approximately 10-fold higher c-di-GMP levels and led to reduced virulence in a subcutaneous infection model (40). In *V. cholerae*, elevated c-di-GMP (visualized by 2D-TLC) led to 5-fold reduced competitive colonization in the infant mouse small intestine model (39). In that study, the authors tested if the reduction was due to overexpression of biofilm (VPS), but it was not, and they attributed the phenotype to the resulting virulence gene expression downstream of c-di-GMP. We note that the magnitude of the effect we observe in *V. fischeri* colonization of bobtail squid is substantially higher: >100-fold in a single-strain colonization assay. It therefore seems likely that this aspect of mutualism is broadly shared with pathogenic colonization, and we expect there to be strong constraints to retain c-di-GMP levels low during the initiation of the host colonization process. There are also likely to be beneficial symbiosis-specific adaptations within c-di-GMP signaling. For example, the zebrafish gut symbiont *Aeromonas veronii* responds to host amino acids to inhibit a diguanylate cyclase, thereby stimulating bacterial motility into the host (41). It remains to be seen whether there are similar specific adaptations in the *V. fischeri*-squid symbiosis.

Despite encoding 50 genes that manipulate c-di-GMP, we were surprised to find that substantially lowering levels of c-di-GMP (approximately 9-fold) (**Fig. 1A**) led to no significant defect in host colonization. In the Low cdG strain, we detected slightly smaller bacterial aggregates in the host mucus (**Fig. 5**), but no consequent diminution of bacterial counts in the light organ at 18 hpi or 48 hpi (**Fig. 2**). The Low cdG strain similarly exhibited no competitive colonization defect (**Fig. S1**). When examining transcriptional reporters for the symbiosis polysaccharide gene *sypA* or the cellulose biosynthesis gene *bcsQ in vivo*, the Low cdG strain resembled the wild-type parent (**Fig. 6A,B, Fig. S6**). We note that colonization was intact despite this strain exhibiting strong biofilm and motility phenotypes *in vitro* (**Fig. 1**), thus emphasizing the importance of studying these phenotypes in the relevant host system. The key bacterial behaviors during the initial stages in the host include adhering to host mucus and then cilia within the mucus (10); forming the Syp-dependent aggregates (5, 9–11); migrating toward the host pores in a manner independent of flagellar motility (4, 12); employing flagellar motility and chemotaxis to swim toward chitin oligosaccharides released by the host (8, 14–17); and resisting host innate immune attacks as they traverse the pore into the duct, antechamber, bottleneck, and then crypts of the light organ (42–45). These processes take 18-24 h and encompass the “initiation” and “accommodation” phases of the symbiosis. Around 18-24 h, the interaction shifts to the “persistence” phase, where quorum sensing stimulates bioluminescence in the host, and a daily (diel) rhythm ensues in which bacteria consume nutrient sources provisioned from the host squid. We observe specific initiation defects in biofilm reporter gene expression, biofilm aggregate number and architecture, flagellar motility, and bacterial counts in the host at 18 hpi. We additionally observe an accommodation defect in that bacterial counts do not recover by 48 hpi, arguing that the high cdG strain continues to inhibit expansion in the crypts following the initial defect. The accommodation phenotype is unlikely to be due to flagellar motility, as most bacteria lose their flagella during the mature association (8). Therefore, these effects may be due to regulation of biofilm genes in the host (7) or may be due to other unrecognized effects in the High cdG strain.

By using strains with altered c-di-GMP levels, we have uncovered regulatory differences in transcriptional reporters in culture versus the host. In culture, we observed a 3.2-fold increase in *bcsQ’-gfp^+^* reporter activity in the WT strain relative to the Low cdG strain (**Fig. 1D**). However, in the animal, we observed similar *bcsQ’-gfp^+^*reporter activity in the two strains (**Fig. 6B, Fig. S6B**), suggesting that the lower *bcsQ’-gfp^+^*activity of the WT strain may occur in response to the host environment. Similarly, the upregulation of *bcs* gene expression in the background of elevated c-di-GMP conditions led to a substantial downregulation of *sypA’-gfp*^+^ reporter activity in the host, yet no comparable effect was observed in two distinct culture conditions (**Fig. 6**). Given the multiple cases in which gene expression in the host is distinct from that observed in culture, it remains an intriguing question as to how the host influences pathways linked to c-di-GMP signaling.

If the Low cdG strain thrives in the host under conditions where levels of c-di-GMP remain low, is c-di-GMP required for *V. fischeri* biology in the squid? Given that there are another twenty-one DGCs and five putative bifunctional DGC/PDEs still present in the Low cdG strain, our results do not preclude a requirement for c-di-GMP during colonization. It remains possible that additional DGCs perform localized functions that were not addressed in the current study and/or that DGCs play critical roles at later stages in the colonization process.

### Construction of multiple-gene deletions in *V. fischeri* is an effective tool to tackle gene families

The recent development of a method for construction of serial deletions (26) enabled the construction of multiple-mutant strains such as the Low cdG and High cdG strains in this study. We have further extended the method to allow for detection of barcoded deletions in competitive colonization experiments (46). The work in this study highlights the utility of such approaches to interrogate gene families. There are many instances in which deletions of multiple genes could clarify the role of various biological processes. Previously, we identified a role for chemotaxis toward chitin oligosaccharides as a key developmental stage during squid colonization (16). Efforts from our group and others have studied 23 *V. fischeri* chemoreceptors through a combination of single and multiple mutants (47–49), but there are a total of 43 methyl-accepting chemoreceptors encoded in the ES114 genome, and the receptor(s) for chitin oligosaccharides remain elusive. Generation of multiple mutant strains has been especially valuable to dissect gene families in *V. cholerae* (50), and our work suggests that a similar approach will be useful in *V. fischeri*.

### Elevated c-di-GMP levels diminish, but do not eliminate, flagellar motility *in vivo*

The High cdG strain has a significant soft agar motility defect (**Fig. 1B**), and amotile strains are at a significant colonization disadvantage compared to WT. Our competition of the High cdG strain with the isogenic Δ*flrA* derivative revealed a 15-fold advantage for the former during squid colonization (**Fig. 4**). Our interpretation of this result is that the High cdG strain retains the ability to be motile in the host despite the elevated c-di-GMP levels. The magnitude of the competitive advantage (15-fold) is diminished compared to that of the wild-type strain competing against a Δ*flrA* strain (>1,000-fold) (46, 51). Therefore, it seems likely that c-di-GMP does impact motility in the host, but not to the extent that it fully prohibits colonization.

### Cyclic di-GMP has a qualitative effect on the nature of the exopolysaccharide in animal tissue

The increased number of aggregates observed in the High cdG strain in squid is, to our knowledge, the first time such a phenotype has been observed (**Fig. 5**). Hypermotile mutants were noted to exhibit more diffuse aggregates on the light organ surface, though these were often smaller than their WT counterparts (15). Multiple studies have noted larger *in vivo* aggregates upon overexpression of biofilm activator RscS or removal of biofilm inhibitor BinK (5, 35). In both cases, it was shown that Syp was the relevant pathway induced, and the cells within the resulting larger aggregates were functional to outcompete those from WT aggregates. The morphology and gene expression in High cdG aggregates are significantly more complicated and implicate an interaction between cellulose and Syp exopolysaccharides. We showed that cellulose on its own is not required for colonization (**Fig. S5B**). However, the high c-di-GMP condition revealed a role for the *bcs* (cellulose synthase) locus in regulating *syp* (mirroring results obtained in specific agar conditions in (52)) (**Fig. 6A, Fig. S6A**). Therefore, while cellulose production on its own does not impact colonization, in the High cdG background it significantly impacts expression of *syp*. In *Pseudomonas aeruginosa*, roles for multiple exopolysaccharide systems in biofilm formation have been explored, for example as the Psl and Pel systems have distinct but also overlapping roles in attachment and biofilm formation (53). In *V. fischeri*, our work supports data in the literature that Syp is the key system for *in vivo* biofilm formation and that the cellulose system is dispensable for host colonization. At the same time, this study reveals a novel transcriptional interaction between two exopolysaccharide systems. This interplay may impact the formation of multiple-exopolysaccharide biofilms during later stages of host colonization or under environmental conditions that have yet to be explored, and points to a novel regulatory mechanism that connects biofilm regulation in an animal host.

In conclusion, this work identified a role for c-di-GMP regulation during the initiation of a mutualism, identified specific developmental stages in the host that were regulated by altered c-di-GMP levels, and revealed a transcriptional interplay between multiple exopolysaccharide systems that is apparent only in the host. Our study therefore lays the groundwork for future investigations into how this widespread bacterial compound influences bacterial behaviors beyond pathogenesis to impact microbiome behavior and interactions with symbiotic hosts.

## MATERIALS AND METHODS

### Bacterial strains, plasmids, and media

*V. fischeri* and *E. coli* strains used in this study are listed in Table 1. Plasmids used in this study are listed in Table 2. *V. fischeri* strains were grown at 25°C in Luria-Bertani salt (LBS) medium (per liter: 25 g Difco LB broth [BD], 10 g NaCl, 50 mL 1 M Tris buffer [pH 7.5]), or TBS (per liter: 10 g tryptone [VWR], 20 g NaCl, 35 mM MgSO_4_ [where noted], 10 mM calcium chloride [where noted], 50 mM 1 M Tris buffer [pH 7.5]) where noted. *E. coli* strains used for cloning and conjugation were grown at 37°C in Luria-Bertani (LB) medium (per liter: 25 g Difco LB broth [BD]). When needed, antibiotics were added to the media at the following concentrations: kanamycin, 100 μg/mL for *V. fischeri* and 50 μg/mL for *E. coli*; chloramphenicol, 1 or 5 μg/mL for *V. fischeri* and 25 μg/mL for *E. coli*; erythromycin, 2.5 or 5 μg/mL for *V. fischeri*; gentamicin, 2.5 μg/mL for *V. fischeri* and 5 μg/mL for *E. coli*; trimethoprim, 10 μg/mL for *V. fischeri*; spectinomycin, 160 μg/mL in LB for *V. fischeri*. When needed, thymidine was added at 0.3 mM for *E. coli*. Growth media was solidified using 1.5% agar when needed. For Congo red agar, 40 μg/mL Congo red and 15 μg/mL Coomassie blue were added to LBS. For Xgal agar, 20 μg/mL Xgal were added to LBS. Plasmids were introduced from *E. coli* strains into *V. fischeri* strains using standard techniques (54, 55).

**Table 1:**
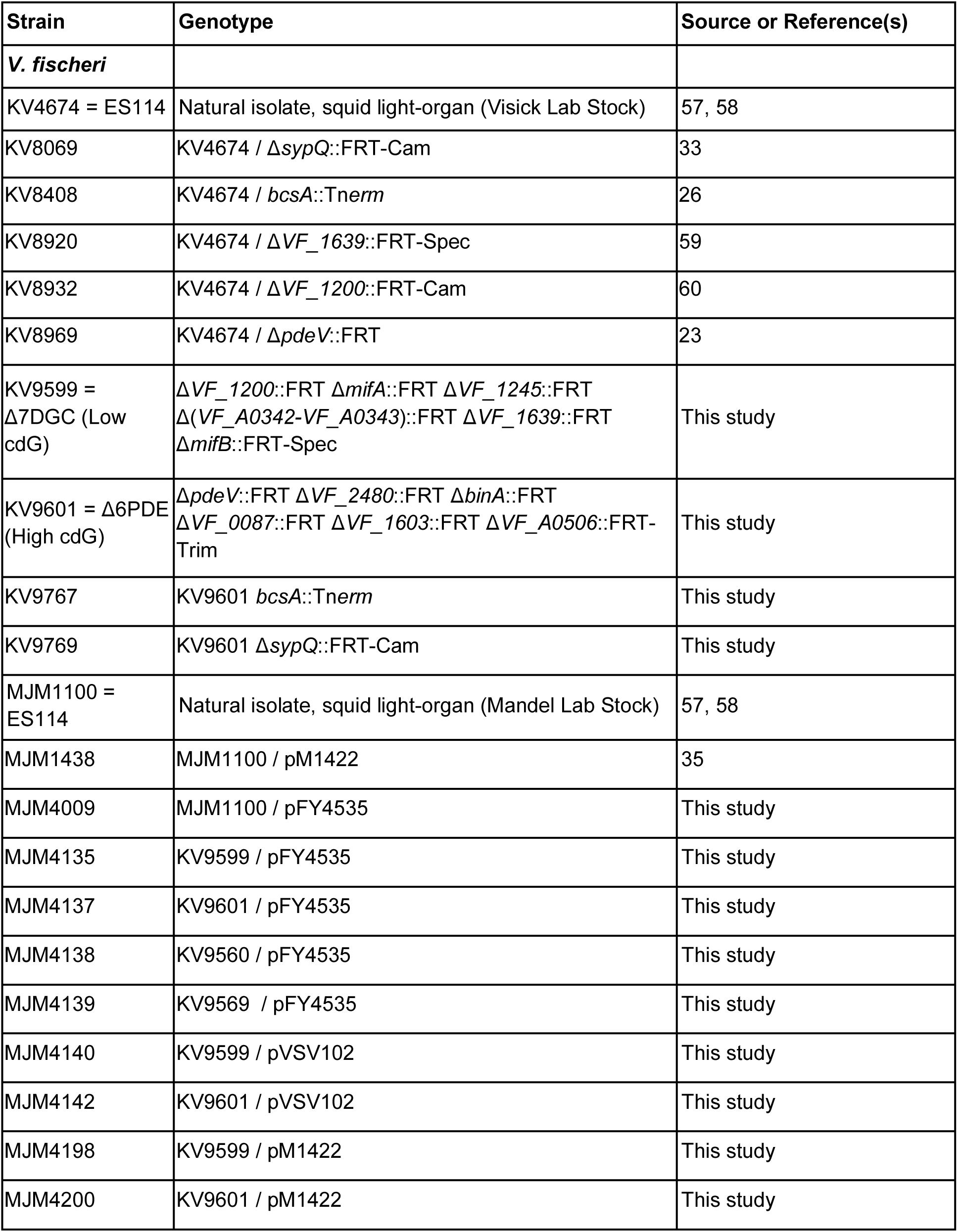
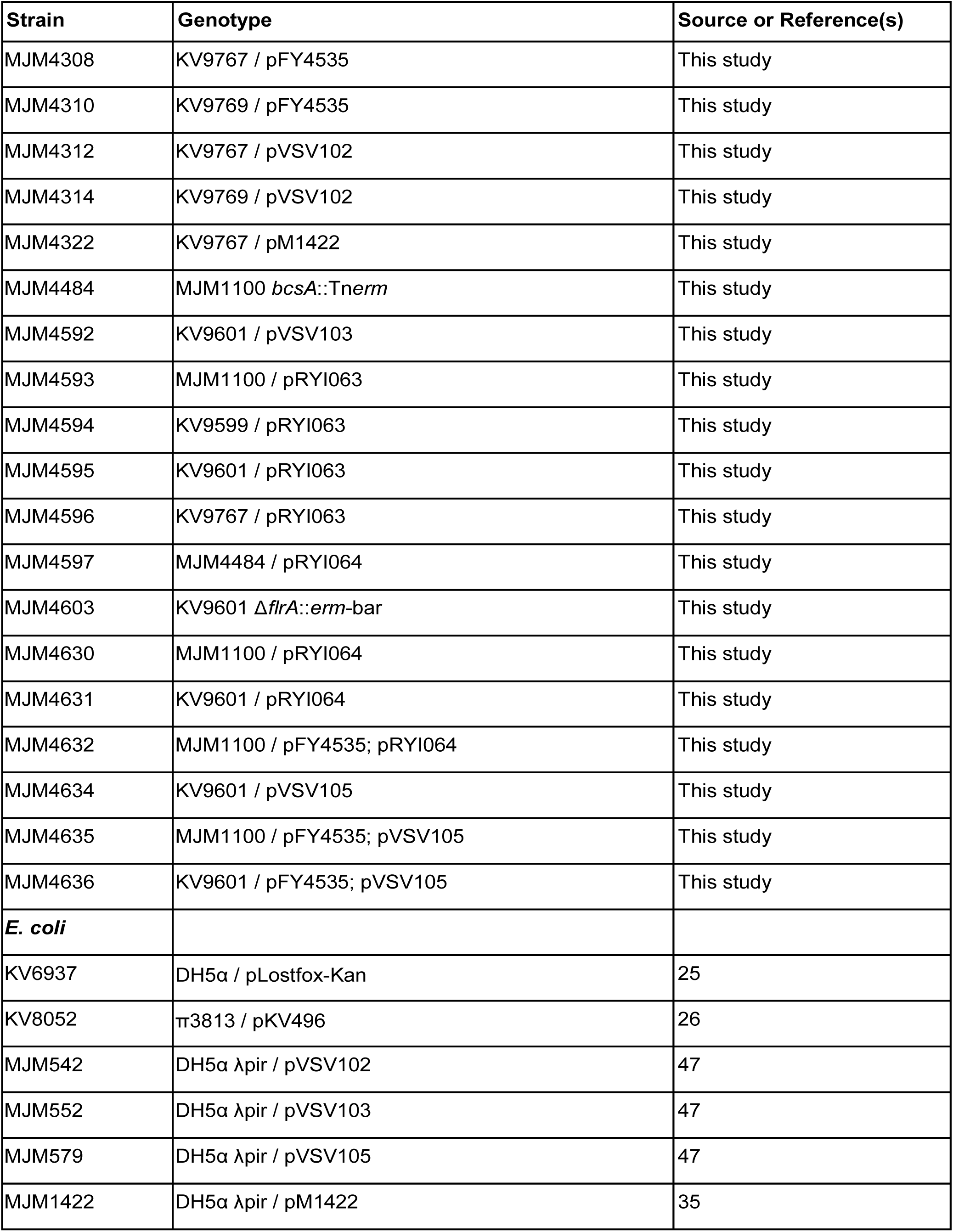
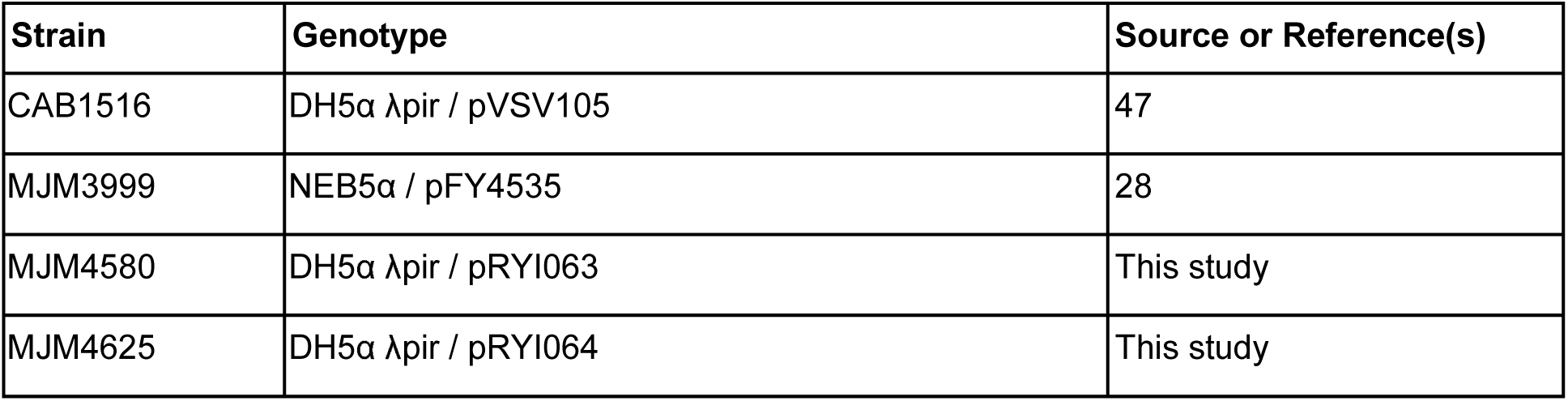
Strains used in this study.

**Table 2:** Plasmids used in this study.

| Plasmid | Description | Source or Reference(s) |
| --- | --- | --- |
| pEVS104 | Conjugal helper plasmid (Kan<sup>R</sup>) | 54 |
| pFY4535 | c-di-GMP reporter plasmid (Gent<sup>R</sup>) | 28 |
| pJJC4 | Vector carrying *tfoX* and *litR* genes (Cam<sup>R</sup>) | 61 |
| pKV494 | Vector carrying FRT-Erm<sup>R</sup> | 26 |
| pKV495 | Vector carrying FRT-Cam<sup>R</sup> | 26 |
| pKV496 | pLostfoX-Kan backbone containing the FLP recombinase (Kan<sup>R</sup>) | 26 |
| pKV521 | Vector carrying FRT-Spec<sup>R</sup> | 26 |
| pLostfoX-Kan | TfoX induction vector for chitin-pathway transformation (Kan<sup>R</sup>) | 25 |
| pM1422 | pTM267 *sypA'*-*gfp*<sup>+</sup> (Cam<sup>R</sup>) | 35 |
| pMCL2 | Vector carrying FRT-Trim<sup>R</sup> | 26 |
| pRYI063 | pTM267 *bcsQ'*-*gfp*<sup>+</sup> (Cam<sup>R</sup>) | This study |
| pRYI064 | pVSV105 carrying *VC1086* (Cam<sup>R</sup>) | This study |
| pVSV102 | Constitutive GFP (Kan<sup>R</sup>) | 62 |
| pVSV103 | Constitutive LacZ (Kan<sup>R</sup>) | 62 |
| pVSV105 | Complementation vector (Cam<sup>R</sup>) | 47 |

### DNA synthesis and sequencing

Primers used in this study are listed in Table 3 and were synthesized by Integrated DNA Technologies (Coralville, IA). Full inserts for cloned constructs and gene deletions were confirmed by Sanger Sequencing at Functional Biosciences via UW-Madison. Sequence data were analyzed using Benchling. PCR to amplify constructs for cloning and sequencing were performed using Q5 High-Fidelity DNA polymerase (NEB), OneTaq DNA polymerase (NEB), or MilliporeSigma Novagen KOD DNA Polymerase. Diagnostic PCR was performed using GoTaq polymerase (Promega) or OneTaq DNA polymerase.

**Table 3:**
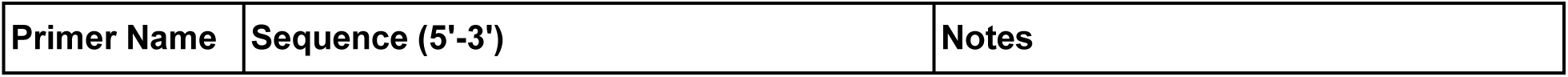
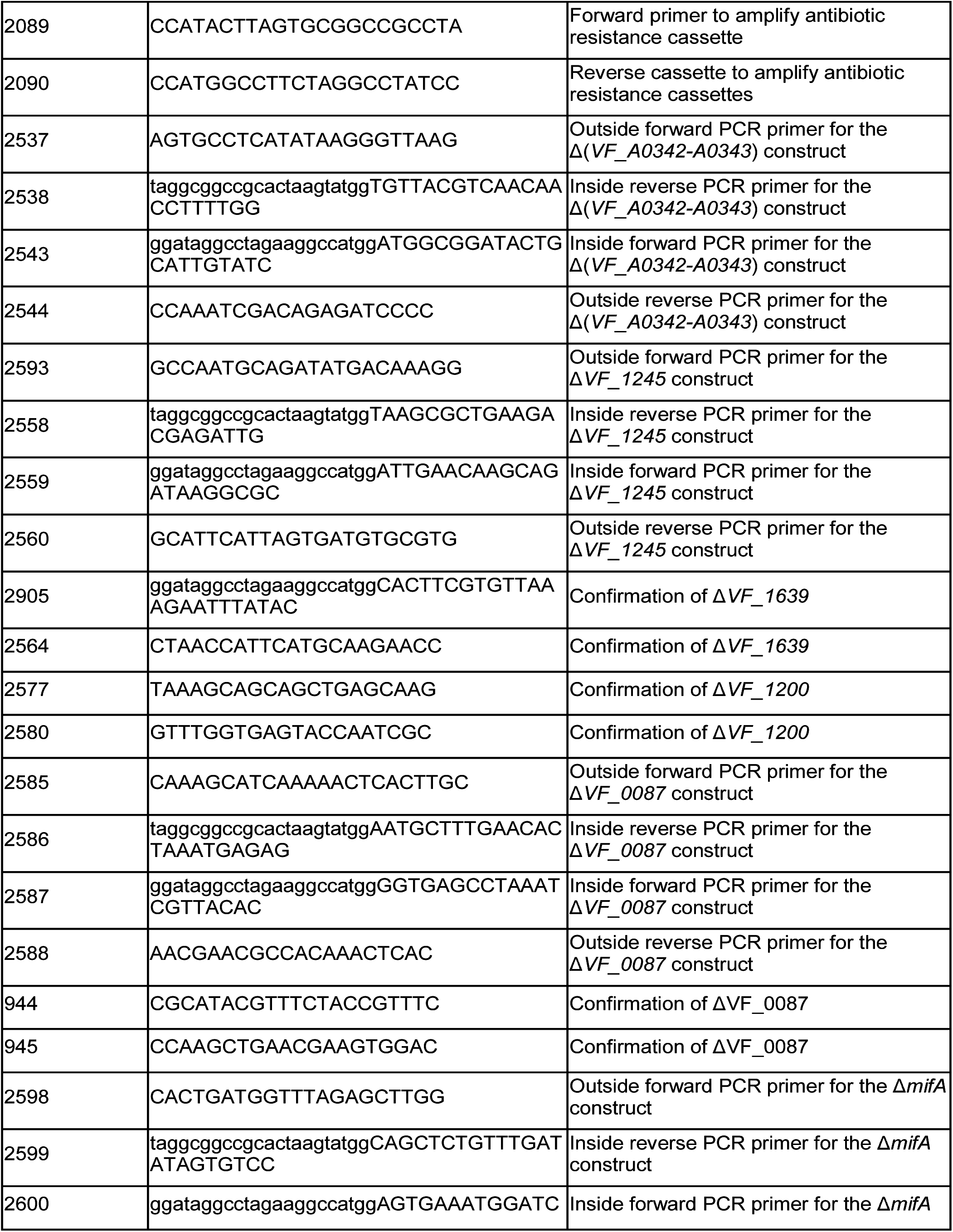
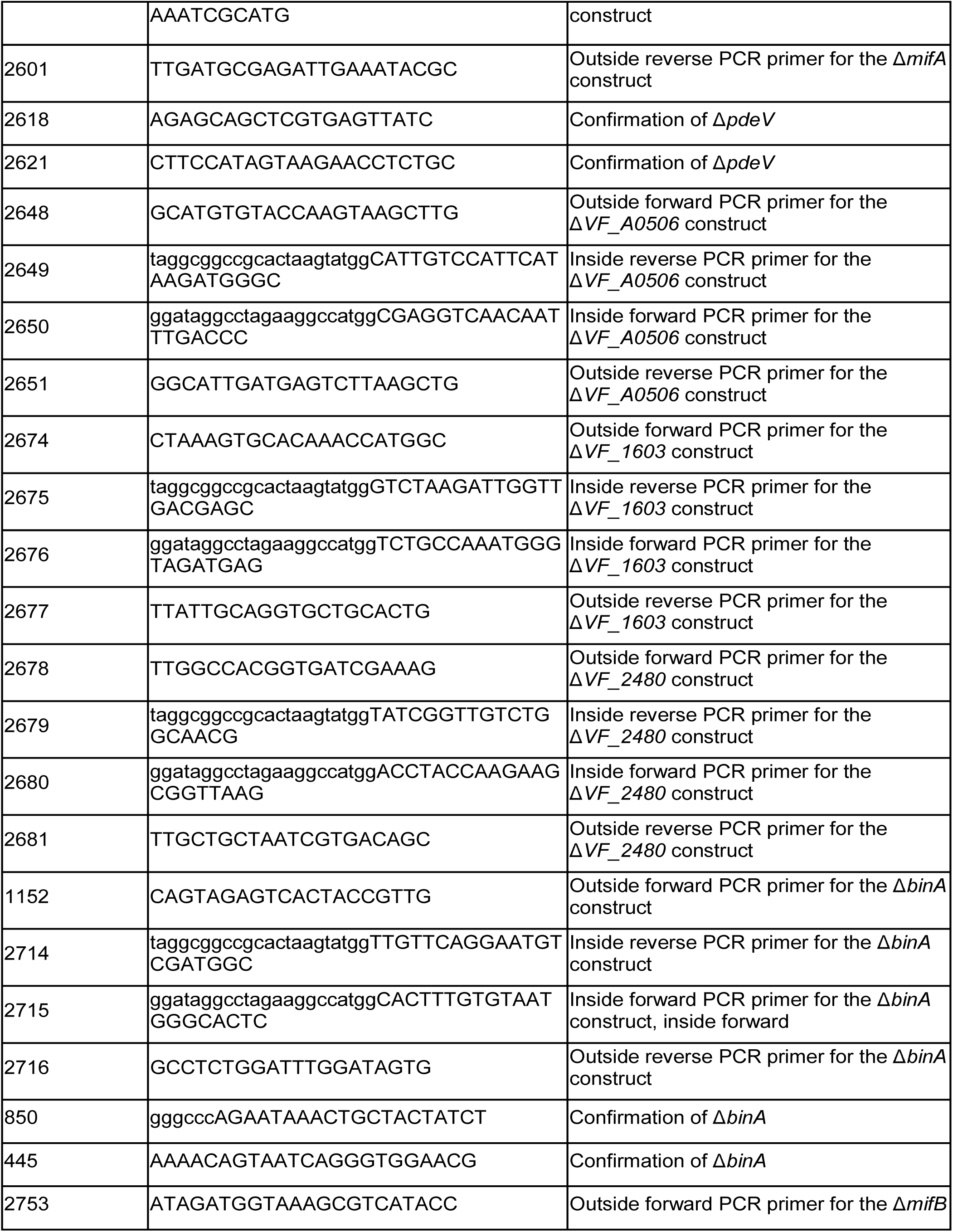
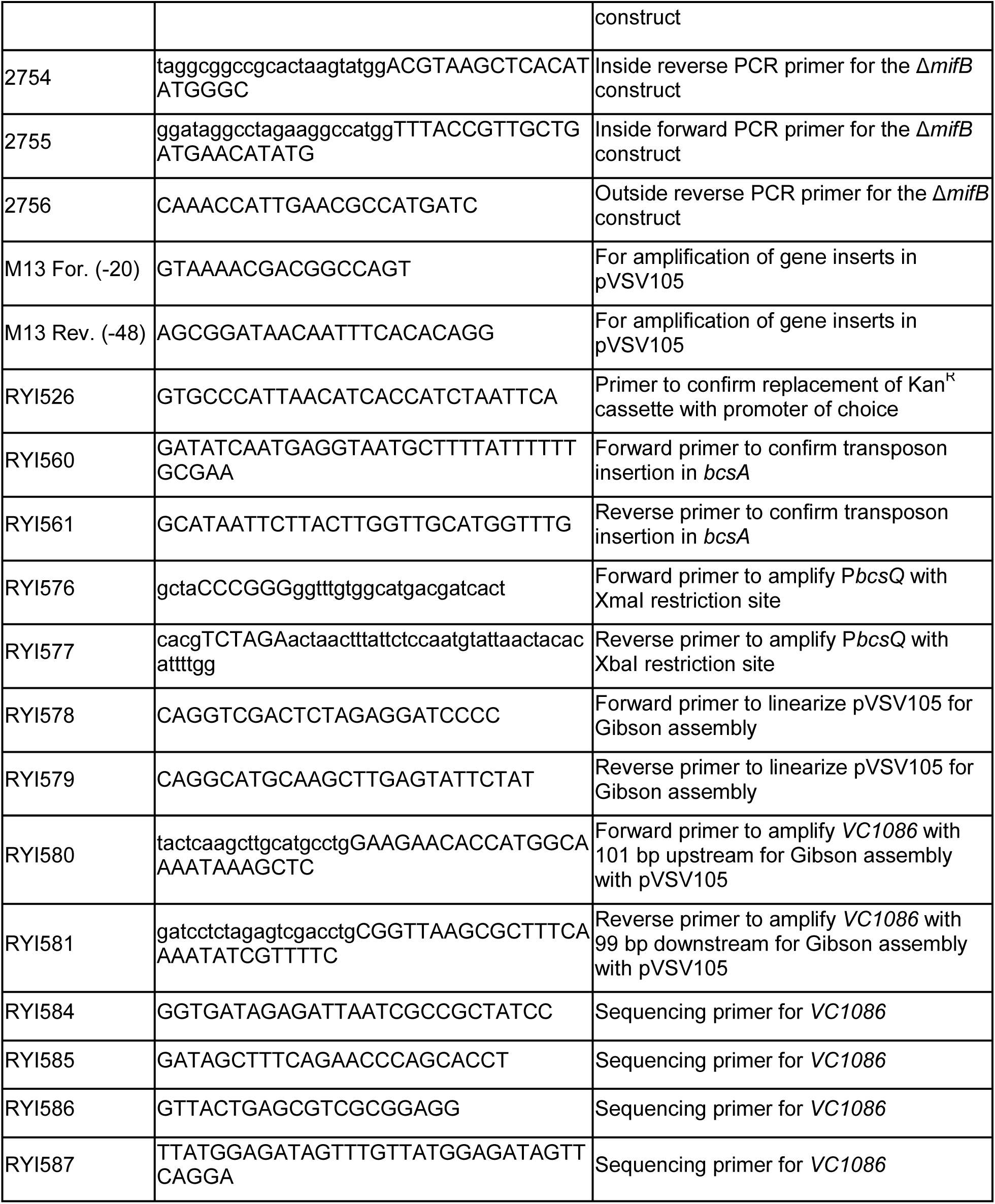
Primer List.

### Construction of multiple gene deletion mutants

The multiple DGC and PDE mutants were constructed as follows. First, single gene deletions were generated using PCR-SOE and *tfoX*-induced transformation methods previously described (26, 56); some of these mutants have been previously described (23, 52). Briefly, regions flanking a gene of interest were amplified using primers listed in Table 3 and joined to an antibiotic-resistance cassette (e.g., Trim^R^/Erm^R^/Spec^R^) flanked by Flp-recombinase target (FRT) sites that was amplified with primers 2089 and 2090. The resulting fused PCR product was transformed into a *V. fischeri* strain (generally ES114) that contained a TfoX-overproducing plasmid (pLostfoX or pLostfoX-Kan plasmid (25, 56). Selection for the integrants was conducted on selective agar. The deletion/insertion was verified by PCR, generally with the outside primers used to generate the flanking fragments. Next, strains with multiple mutations were generated using genomic DNA from specific single mutants as follows. A *tfoX*-overexpression plasmid was introduced into a mutant of interest and transformed with genomic DNA containing a specific mutation, followed by selection for and verification of the mutation as described above. The resulting strain carrying a *tfoX*-overexpression plasmid could then serve as a recipient for the introduction of the next mutation. Because some mutants were generated using the same antibiotic resistance cassette, as needed the Flp-encoding plasmid pKV496 was introduced to induce the loss of the antibiotic resistance cassette and the retention of a 112 bp FRT scar in the gene. Following strain construction, the integrity of each deletion junction was verified with diagnostic primers listed in Table 3 to ensure that only recombination at local FRT sites occurred. As noted in the strain list (Table 1), the final cassette often was left in place. As an example, for Δ7PDE strain KV9601, the starting strain, KV8969 (23) (Δ*pdeV*::FRT) carrying *tfoX* plasmid pLostfoX-Kan was sequentially transformed with genomic DNA carrying the Δ*VF_2480*::FRT-Erm, Δ*binA*::FRT-Trim, and Δ*VF_0087*::FRT-Spec mutations. The cassettes were removed, and the *tfoX* plasmid reintroduced. Next, the Δ*VF_1603*::FRT-Trim mutation was introduced, and the cassette removed. Finally, the Δ*VF_A0506*::FRT-Trim mutation was introduced. Similar methods were used to generate KV9599 (Δ7DGC) and the derivatives of KV9601 that carry mutations in *sypQ* or *bcsA* (the latter of which was derived from strain KV8408, which carries a Tn mutation in *bcsA* (26)) rather than a cassette as described above).

### Construction of *bcsA*::Tn*erm* strain

Genomic DNA isolated from KV9767 (Δ6PDE *bcsA*::Tn*erm*) was introduced into MJM1100 via transformation using pLostfoX-Kan (25, 56). Mutant candidates were selected using erythromycin and screened by PCR using RYI560 and RYI561 primers.

### Construction of the *bcsQ*’-*gfp*^+^ transcriptional reporter plasmid pRYI063

The *bcsQ* promoter (*bcsQ* -444 to -29; P*bcsQ*) was amplified from MJM1100 genomic DNA using RYI576 and RYI577 primers that added flanking XmaI and XbaI restriction sites. The amplified P*bcsQ* and pTM267 were digested with XmaI and XbaI. Digested pTM267 was treated with antarctic phosphatase. Digested P*bcsQ* was ligated into digested pTM267 using T4 DNA ligase (NEB). The ligation reaction was transformed into chemically competent DH5α λpir. Transformant candidates were selected using chloramphenicol and confirmed by Sanger sequencing using the RYI526 primer.

### Construction of the *VC1086* expression plasmid pRYI064

pVSV105 was amplified using RYI578 and RYI579 primers. *VC1086*, along with 101 bp upstream and 99 bp downstream, was amplified from *Vibrio cholerae* C6706 str2 genomic DNA using RYI580 and RYI581 primers. *VC1086* was assembled with pVSV105 by Gibson assembly using NEBuilder HiFi DNA Assembly Master Mix (NEB). The Gibson reaction was transformed into chemically competent DH5α λpir and transformant candidates were selected using chloramphenicol. Plasmid was screened by PCR using M13 For. (-20) and M13 Rev. (-48) primers. Plasmid was confirmed by Sanger sequencing using M13 For. (-20), M13 Rev. (-48), RYI584, RYI587, RYI585, and RYI586 primers.

### C-di-GMP quantification

Strains were streaked on LBS agar and incubated overnight at 25°C. Liquid LBS was inoculated from single colonies and grown at 25°C overnight on a rotator. 8 μL of liquid culture were spotted onto LBS agar and incubated at 25°C for 24 h. Spots were resuspended in 1 mL 70% Instant Ocean (IO). OD_600_, TurboRFP (555 nm excitation/585 nm emission), and AmCyan (453 nm excitation/486 nm emission) for each resuspended spot were measured in triplicate using BioTek Synergy Neo2 plate reader. To calculate c-di-GMP levels, TurboRFP values (reports c-di-GMP levels) were normalized to AmCyan values (constitutively expressed).To compare c-di-GMP levels of strains in LBS vs. IO, 1 mL of overnight LBS liquid cultures were pelleted at 8000 x *g* for 10 min, resuspended in 600 μL filter-sterilized IO (FSIO), and both the FSIO and LBS samples were incubated at room temperature for 3 hr; LBS and FSIO strains were pelleted at 8000 x *g* for 10 min, and resuspended in 600 μL FSIO. OD_600_, TurboRFP, and AmCyan for each sample were measured in triplicate as described above.

### Motility assay

Strains were streaked on TBS and TBS-Mg^2+^ agar and incubated overnight at 25°C. Single colonies were inoculated into TBS and TBS-Mg^2+^ soft agar (0.3% agar), respectively, and incubated at 25°C for 5 h. Images of plates were taken using a Nikon D810 digital camera and diameter of migration was measured using ImageJ software.

### Isolation of cells from the outer edge of motility ring

Strain was streaked on TBS-Mg^2+^ agar and incubated overnight at 25°C. A single colony was inoculated into TBS-Mg^2+^ soft agar (0.3% agar) and incubated at 25°C for 18 h. Cells from the outer edge of the resulting motility ring were streak purified on LBS. Isolates were then reinoculated into new TBS-Mg^2+^ motility agar as described above.

### Congo red assay

Strains were streaked on LBS agar and incubated overnight at 25°C. Liquid LBS was inoculated with single colonies and grown at 25°C overnight on a shaker. 4 μL spots of liquid culture were spotted on LBS Congo red agar and incubated 24 h at 25°C. Spots were transferred onto white printer paper (26) and images were scanned as TIFF files. Congo red binding was quantified using ImageJ software.

### *syp* and *bcs* transcriptional reporter analysis *in vitro*

Strains were streaked on LBS agar and incubated overnight at 25°C. Liquid media was inoculated with single colonies and grown at 25°C for 24 hr on a shaker. 4 μL of overnight cultures were spotted onto TBS or TBS-Ca^2+^ agar plates and grown at 25°C for 24 hr (52). Spots were imaged using the Zeiss Axio Zoom.v16 large-field 561 fluorescent stereo microscope and fluorescence levels were measured using the Zen Blue Software polygon tool. To calculate gene promoter activity, GFP values (reports promoter activity) were normalized to mCherry values (constitutively expressed).

### Growth curves

Strains were streaked on LBS agar and incubated overnight at 25°C. Liquid LBS was inoculated with single colonies of each strain in triplicate and grown at 25°C overnight on a rotator. Cultures were diluted 1:1000 in LBS and grown at 25°C on a rotator to OD_600_ ∼0.5. Cultures were diluted 1:100 in LBS (n = 6 technical replicates per culture) in a 96-well microtiter plate. OD_600_ was measured every 15 min for 20 hr at 25°C using BioTek Synergy Neo2 plate reader. Doubling time was calculated in Prism using a nonlinear regression analysis of each strain during exponential phase.

### Squid single strain colonization assay

*E. scolopes* hatchlings were colonized with approximately 10^3^-10^5^ CFU/mL of bacteria according to standard procedure (29). At 18 hpi and 48 hpi, luminescence of hatchlings was measured using the Promega GloMax 20/20 luminometer and euthanized by storage at -80°C. CFU counts per light organ were determined by plating homogenized euthanized hatchlings and counting colonies.

### Squid competitive colonization assay

Strains were grown overnight in liquid LBS at 25°C on a rotator. Strains were diluted 1:80 in liquid LBS and grown to an OD_600_ of approximately 0.2. The strains were mixed in a 1:1 ratio with a competing strain carrying a constitutive *lacZ* on the pVSV103 plasmid and the mixed cultures were used to inoculate *E. scolopes* hatchlings with approximately 10^3^-10^4^ CFU/mL. At 3 hpi, hatchlings were washed and transferred to 40 mL filter-sterilized Instant Ocean (FSIO). Water was changed at 24 hpi and hatchlings were euthanized at 48 hpi by storage at -80°C. Euthanized hatchlings were homogenized and plated on LBS-Xgal agar. Competitive index of strains was measured by calculating the blue/white colony ratios as previously described (25, 29).

### Squid aggregation assays

Strains were grown overnight in liquid LBS at 25°C on a rotator. *E. scolopes* hatchlings were inoculated with approximately 10^5^-10^8^ CFU/mL of bacteria. At 3 hpi, squid were anesthetized in 2% EtOH in FSIO and immediately dissected and imaged as described below, or were fixed in 4% paraformaldehyde in 1x mPBS (50 mM phosphate buffer, 0.45 M NaCl, pH 7.4) for approximately 48 h. Fixed hatchlings were washed in 1x mPBS x4, dissected, and imaged using the Zeiss Axio Zoom.v16 large-field 561 fluorescent stereo microscope. Aggregate area was selected and fluorescence levels were measured using the Zen Blue Software polygon tool. Note that the analyses were conducted with all data as well as solely with the inocula in the 10^6^-10^7^ range and the same results were observed, so we included all of the data in the manuscript.

### Data analysis

Congo red binding was quantified using ImageJ by subtracting the WT gray value from the mutant gray value and multiplying the value by -1. Fluorescence of strains in liquid culture were measured using a BioTek Synergy Neo2 plate reader. Fluorescence of strains spotted on agar plates and bacterial aggregates in the squid were measured using Zen Blue Software.

GraphPad Prism was used to generate graphs and conduct statistical analyses. Graphs were further refined in Adobe Illustrator.

## ACKNOWLEDGMENTS

We thank Fitnat Yildiz for providing the pFY4535 reporter plasmid, Chris Waters for input on this project, and Ali Razvi and Steven Eichinger for help with strain construction. This work was funded by NIGMS grant R35 GM119627 to M.J.M., NIGMS grant R35 GM130355 to K.L.V, and R.Y.I. was supported by NIGMS training grant T32 GM007215.

---

**FIG S1** Low levels of c-di-GMP do not inhibit host colonization. Quantification of squid competitive colonization index at 48 hpi by indicated *V. fischeri* strains. Competitive index represents log_10_((test strain/WT)_output_ / (test strain/WT)_input_). Box-and-whisker plots represent minimum, 25th percentile, median, 75th percentile, and maximum. Sample sizes from left to right are 36, 26, and 20 squid. Kruskal-Wallis test was performed for statistical analysis; ns = sot significant, ****p < 0.0001.

**FIG S2** The High c-di-GMP *V. fischeri* strain does not have a growth defect. (A) Growth curves for *V. fischeri* and indicated mutants. For each strain, *n* = 3 biological and *n* = 6 technical replicates per biological replicate. Points represent the mean of technical replicates. Error bars represent standard error of the mean. (B) Doubling times for *V. fischeri* and indicated mutants. Average bars represent the mean of biological replicates. Points represent the mean of technical replicates. Error bars represent standard error of the mean. One-way ANOVA was used for statistical analysis; ns = not significant. Doubling time was calculated using a nonlinear regression analysis of each strain during exponential growth phase. For panels A and B, outliers were excluded from analysis.

**FIG S3** High cdG motile cells are not suppressor mutants. Representative image of migration through soft (0.3%) agar for *V. fischeri* and indicated strains.

**FIG S4** Multiple gene deletions alter *in vivo* c-di-GMP levels. (A) Quantification of c-di-GMP levels for indicated *V. fischeri* strains using the pFY4535 c-di-GMP reporter plasmid in aggregates within the host mucus. Samples sizes from left to right are 9 and 5 aggregates. A Mann-Whitney test was used for statistical analysis; ***p ≤ 0.01. (B) Representative fluorescent microscopy images of squid light organs containing indicated *V. fischeri* strains carrying the pFY4535 c-di-GMP reporter plasmid. Arrows indicate location of aggregates.

**FIG S5** Absence of the cellulose synthase BcsA does not rescue the high c-di-GMP strain colonization defect. (A) Quantification of squid colonization levels at 18 hpi by indicated *V. fischeri* strains and aposymbiotic (Apo) control. Sample sizes from left to right are 14, 25, 17, and 13 squid. (B) Quantification of squid colonization levels at 48 hpi by indicated *V. fischeri* strains. Sample sizes from left to right are 68, 16, 75, 56, 17, and 65 squid. For panels A and B, data for WT, High cdG, and Apo groups are the same as in Fig 3. Box-and-whisker plots represent minimum, 25th percentile, median, 75th percentile, and maximum. Kruskal-Wallis test was performed for statistical analysis for squid that were introduced to bacteria; ns = not significant, **p = 0.002, ****p < 0.0001.

**FIG S6** High c-di-GMP-dependent Bcs expression inhibits *syp* expression in bacterial aggregates within the host mucus. (A) Representative fluorescent microscopy images of squid light organs containing indicated *V. fischeri* strains carrying the pM1422 *sypA*’-*gfp*^+^ transcriptional reporter plasmid. (B) Representative fluorescent microscopy images of squid light organs containing indicated *V. fischeri* strains carrying the pRYI063 *bcsQ*’-*gfp*^+^ transcriptional reporter plasmid.

