## Supplementary material for "High levels of cyclic diguanylate interfere with beneficial bacterial colonization": Figure S1

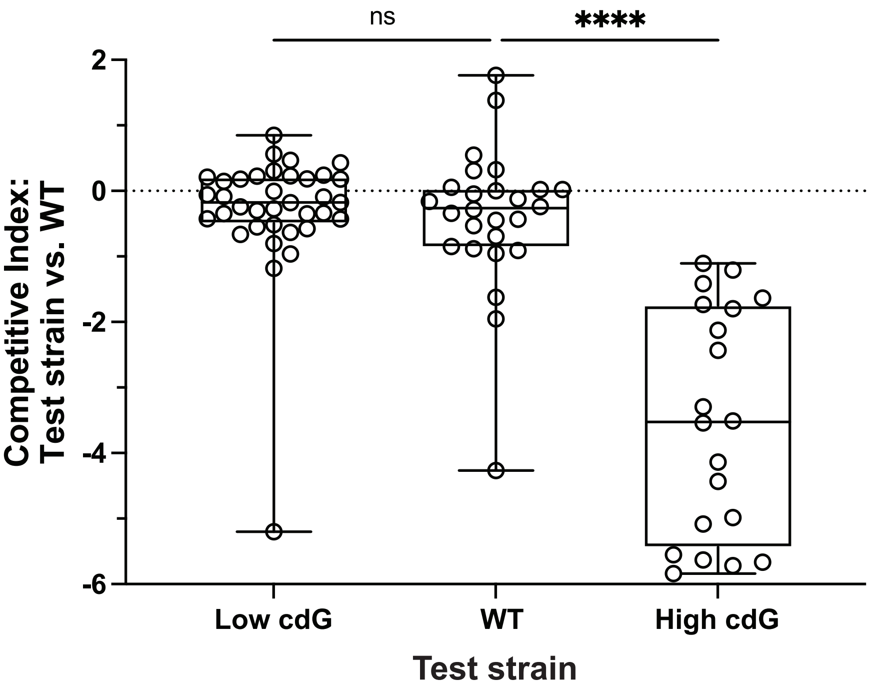


**FIG S1** Low levels of c-di-GMP do not inhibit host colonization. Quantification of squid competitive colonization index at 48 hpi by indicated *V. fischeri* strains. Competitive index represents $\log_{10}$((test strain/WT)$_{output}$ / (test strain/WT)$_{input}$). Box-and-whisker plots represent minimum, 25th percentile, median, 75th percentile, and maximum. Sample sizes from left to right are 36, 26, and 20 squid. Kruskal-Wallis test was performed for statistical analysis; ns = not significant, ****p < 0.0001.
