## Supplementary material for "High levels of cyclic diguanylate interfere with beneficial bacterial colonization": Figure S2

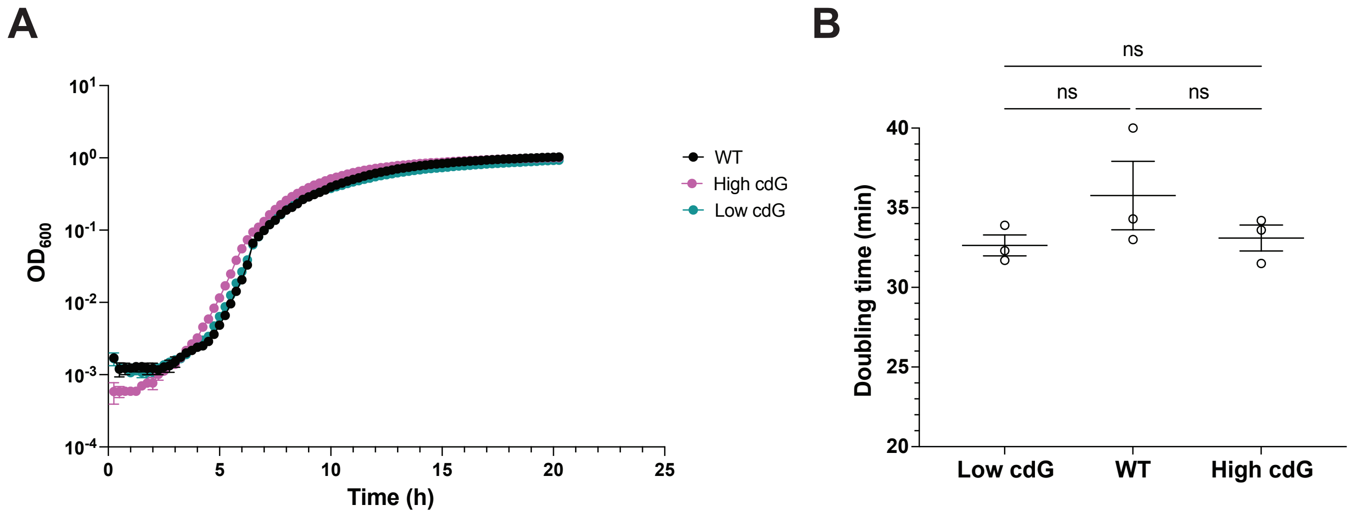


**FIG S2** The High c-di-GMP *V. fischeri* strain does not have a growth defect. (A) Growth curves for *V. fischeri* and indicated mutants. For each strain, $n$ = 3 biological and $n$ = 6 technical replicates per biological replicate. Points represent the mean of technical replicates. Error bars represent standard error of the mean. (B) Doubling times for *V. fischeri* and indicated mutants. Average bars represent the mean of biological replicates. Points represent the mean of technical replicates. Error bars represent standard error of the mean. One-way ANOVA was used for statistical analysis; ns = not significant. Doubling time was calculated using a nonlinear regression analysis of each strain during exponential growth phase. For panels A and B, outliers were excluded from analysis.
