## Supplementary material for "High levels of cyclic diguanylate interfere with beneficial bacterial colonization": Figure S3

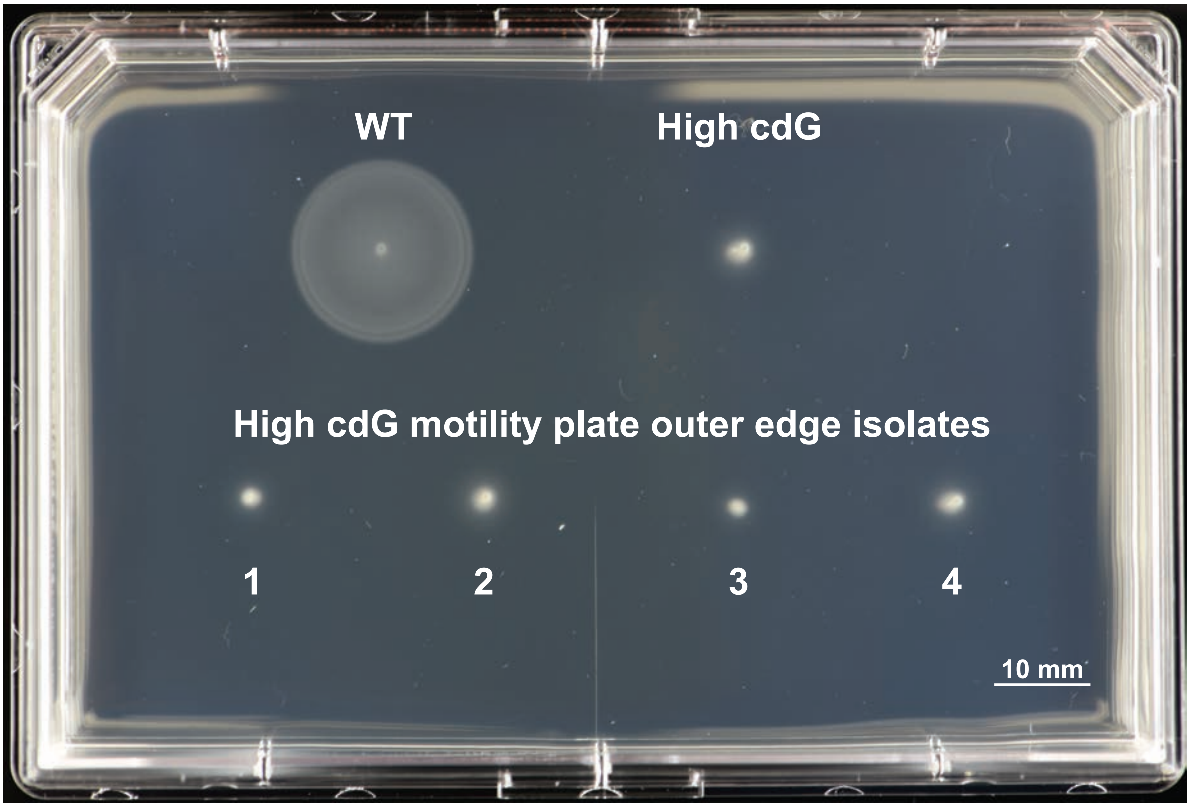


**FIG S3** High cdG motile cells are not suppressor mutants. Representative image of migration through soft (0.3%) agar for *V. fischeri* and indicated strains.
