## Supplementary material for "High levels of cyclic diguanylate interfere with beneficial bacterial colonization": Figure S4

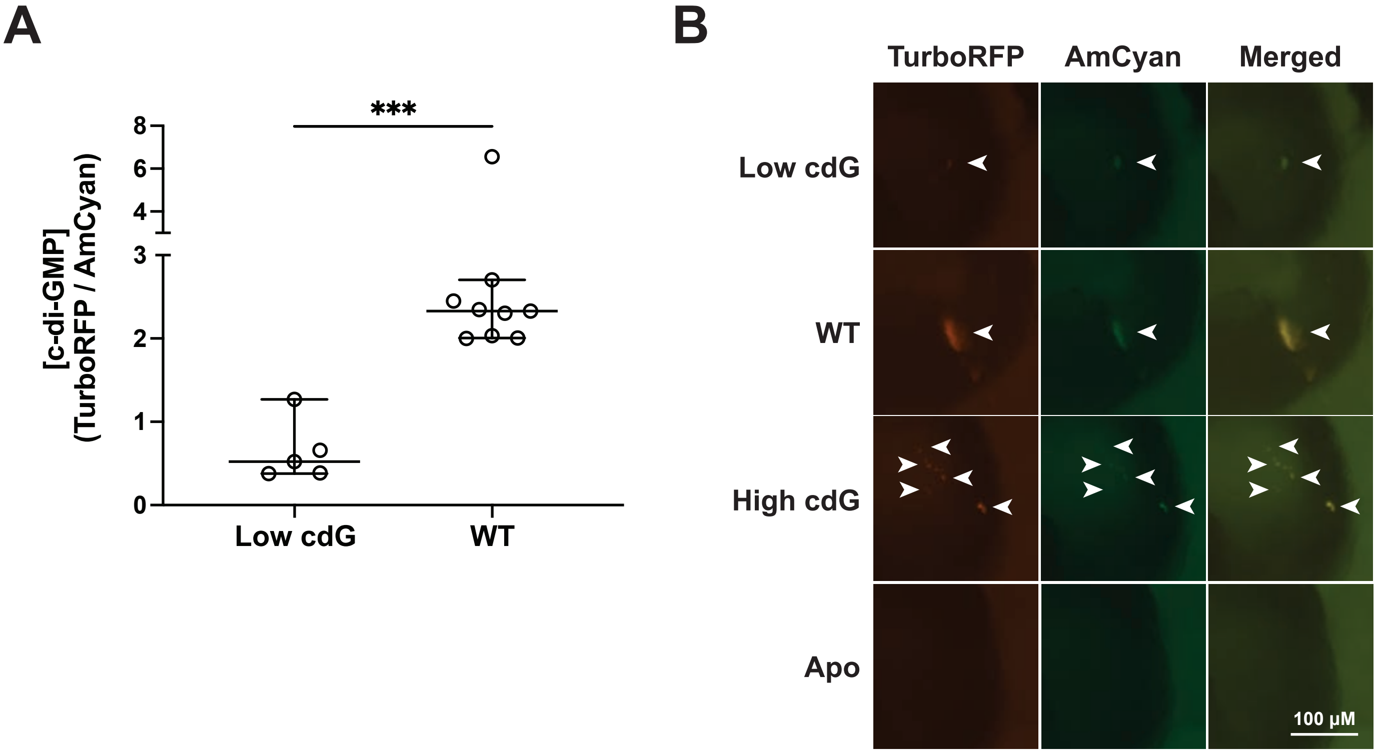


**FIG S4** Multiple gene deletions alter *in vivo* c-di-GMP levels. (A) Quantification of c-di-GMP levels for indicated *V. fischeri* strains using the pFY4535 c-di-GMP reporter plasmid in aggregates within the host mucus. Samples sizes from left to right are 9 and 5 aggregates. A Mann-Whitney test was used for statistical analysis; ***$p \leq 0.01$. (B) Representative fluorescent microscopy images of squid light organs containing indicated *V. fischeri* strains carrying the pFY4535 c-di-GMP reporter plasmid. Arrows indicate location of aggregates.
