## Supplementary material for "High levels of cyclic diguanylate interfere with beneficial bacterial colonization": Figure S5

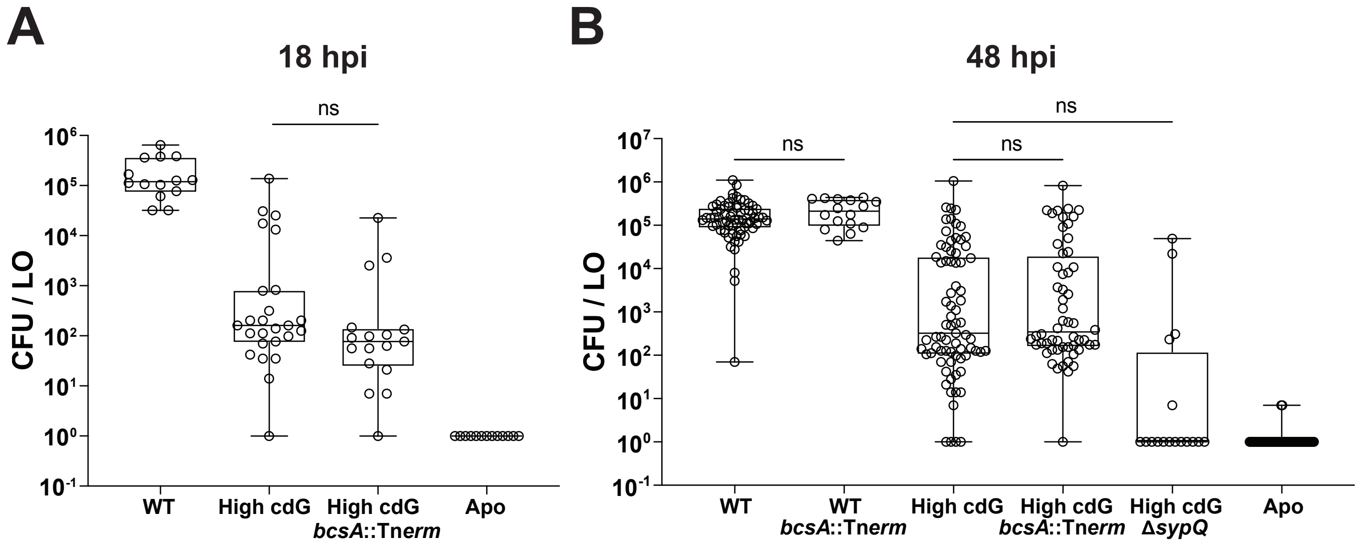


**FIG S5** Absence of the cellulose synthase BcsA does not rescue the high c-di-GMP strain colonization defect. (A) Quantification of squid colonization levels at 18 hpi by indicated *V. fischeri* strains and aposymbiotic (Apo) control. Sample sizes from left to right are 14, 25, 17, and 13 squid. (B) Quantification of squid colonization levels at 48 hpi by indicated *V. fischeri* strains. Sample sizes from left to right are 68, 16, 75, 56, 17, and 65 squid. For panels A and B, data for WT, High cdG, and Apo groups are the same as in Fig 3. Box-and-whisker plots represent minimum, 25th percentile, median, 75th percentile, and maximum. Kruskal-Wallis test was performed for statistical analysis for squid that were introduced to bacteria; ns = not significant, **p = 0.002, ****p < 0.0001.
