## Supplementary material for "High levels of cyclic diguanylate interfere with beneficial bacterial colonization": Figure S6

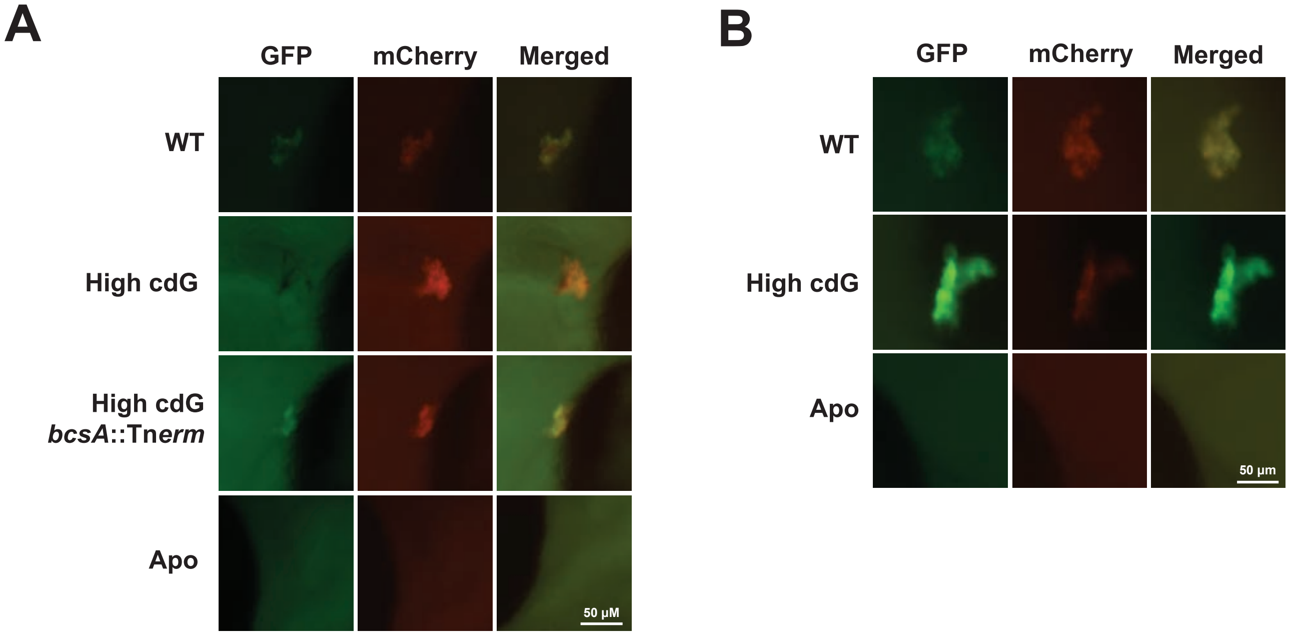


**FIG S6** High c-di-GMP-dependent Bcs expression inhibits *syp* expression in bacterial aggregates within the host mucus. (A) Representative fluorescent microscopy images of squid light organs containing indicated *V. fischeri* strains carrying the pM1422 *sypA*'-*gfp*$^+$ transcriptional reporter plasmid. (A) Representative fluorescent microscopy images of squid light organs containing indicated *V. fischeri* strains carrying the pRYI063 *bcsQ*'-*gfp*$^+$ transcriptional reporter plasmid.
